# Temporal modulation of cortical parvalbumin and somatostatin neurons shapes network activation and adaptation during microstimulation

**DOI:** 10.64898/2026.10.17.759837

**Authors:** Christopher Hughes, Takashi D. Y. Kozai

**Author notes:** Correspondence: C. Hughes or T.D.Y. Kozai.

## Abstract

Parvalbumin-positive (PV) and somatostatin-positive (SOM) inhibitory interneurons play distinct and critical roles in cortical sensory processing, yet their activation dynamics during patterned input remain unclear. This study used two-photon calcium imaging to measure how PV and SOM interneurons are activated and regulate layer 2/3 somatosensory cortical activity during prolonged, patterned input with 22 clinically relevant microstimulation patterns divided by temporal pattern (fixed, burst, or dynamic/biomimetic) and amplitude. Activation over time depended on the interaction between microstimulation amplitude, frequency, and duration. For fixed-frequency stimulation, PV neurons were generally more strongly activated than unlabeled neurons, while SOM neurons were more strongly engaged at higher amplitudes and frequencies, corresponding with greater suppression of surrounding populations over time. Burst patterns limited suppression by evoking stronger and more sustained activation than fixed-frequency patterns especially in PV neurons, potentially by reducing SOM feedback. Theta-burst stimulation (TBS) strongly activated and facilitated SOM neurons, which may underlie its therapeutic effects. Dynamic amplitude modulation preserved excitatory-inhibitory balance by eliciting brief, consistent responses that resemble natural sensory input while reducing long-term adaptation. Many findings fit a model in which PV provides fast-adapting feedforward inhibition and SOM provides slower-facilitating feedback. Supporting this model, distant PV neurons reached peak activation earlier than non-PV neurons, and SOM neurons later than non-SOM neurons during stimulation bursts. A simple computational model built on these dynamics reproduced observed responses, including distance-related effects. Neurons recruited directly bypass SOM input onto their dendrites leading to rapid PV-mediated depression but increasing activity across the stimulation pattern. Together, these results inform the design of microstimulation-based sensory restoration and clarify the circuit-level roles of PV and SOM neurons in transient and sustained inhibitory control of somatosensory cortex.

## Introduction

Microstimulation of the somatosensory cortex can restore tactile sensations in individuals who have lost sensory function^1–5^. In addition to its translational importance, microstimulation provides a powerful experimental approach for probing how cortical circuits respond to temporally patterned input^6–8^. Animals can discriminate microstimulation frequencies in a manner similar to natural tactile stimuli^9–11^, and stimulation frequency systematically alters perceptual quality and intensity in an electrode-dependent manner in humans^5^. Temporal patterns therefore recruit cortical networks in different ways. However, the circuit-level mechanisms behind these effects remain poorly understood, particularly because direct measurements of neural activity during microstimulation remain difficult in humans^12^.

Two-photon calcium imaging in transgenic mouse models has provided new insight into the spatiotemporal dynamics evoked by microstimulation^13–21^. Stimulation frequency strongly shapes spatiotemporal recruitment of cortical activity. High-frequency stimulation rapidly adapts activity in neurons distant from the electrode, while lower frequencies maintain recruitment over prolonged stimulation periods^13,20^. Individual neurons exhibit diverse temporal tuning profiles, with some neurons responding preferentially to low frequencies, high frequencies, or burst-like patterns^14,22^. The frequency reported as most effective differs across studies, alongside differences in cortical area, anesthetic state, stimulus duration, and stimulus amplitude^13–21^. Non-neuronal cells respond on these same timescales. For example, microglial processes orient toward stimulated neurons, and pericytes relax again as stimulation frequency rises, indicating that non-neuronal cells can influence activation and adaptation of neurons^23,24^. Together, these findings indicate that individual cortical neurons as well as non-neuronal cells, differentially encode temporal features of electrical stimulation and suggest that microstimulation provides a framework for studying how cortical circuits process patterned input.

Inhibitory interneurons are likely central to these dynamics given their critical role in sensory processing^25–28^. Two recent studies examining inhibitory neuron responses to microstimulation both found that inhibitory neurons are often more strongly activated than excitatory neurons, with distance and duration dependence^17,19^. Electrical stimulation can also suppress activity in excitatory and inhibitory neurons, shown with voltage-imaging and electrophysiology^29–31^. Previous slice work delivering 10- and 40-Hz stimulation documented suppression of activity especially at 40 Hz, abolished by GABA_A_ antagonists^29^. Prolonged stimulation can also alter excitatory-inhibitory balance, with theta-burst stimulation (TBS) producing strong recruitment of inhibitory activity^17^. These observations support the hypothesis that inhibitory circuits contribute to the suppression or adaptation of cortical responses during sustained stimulation^12^. However, relatively little is known about how inhibitory subtypes respond to different temporal stimulation patterns or how these responses shape broader network recruitment.

Two major inhibitory subclasses, parvalbumin-expressing (PV) and somatostatin-expressing (SOM) interneurons, have been extensively studied in sensory cortex and play distinct, and often opposing, roles in cortical computation and sensory processing^28,32–37^. PV neurons are generally associated with rapid feedforward inhibition and precise temporal control of excitatory neuron spiking, while SOM interneurons are more commonly linked to feedback inhibition, dendritic modulation, and disinhibitory circuit motifs through inhibition of PV neurons^27,28,33^. Both subtypes also exhibit distinct intrinsic electrophysiological properties, with PV neurons often identified as fast-spiking, while SOM neurons more commonly display slower-spiking or bursting dynamics^27,33–35,38,39^. These differences suggest that inhibitory subtypes may entrain on different timescales to different temporal patterns of stimulation and therefore contribute differently to cortical responses during microstimulation.

Here, we investigated how PV and SOM inhibitory neuron subtypes are recruited by different microstimulation patterns in the mouse somatosensory cortex using two-photon calcium imaging. We examined how distinct temporal stimulation patterns influenced the activation dynamics of these inhibitory populations and compared their responses to those of the surrounding cortical network. By characterizing subtype-specific recruitment during microstimulation, this work clarifies how inhibitory circuitry shapes cortical responses to patterned input. More broadly, these experiments provide insight into the functional response properties of PV and SOM neurons and how distinct inhibitory subnetworks process temporally structured activity in sensory cortex. The present work informs both cortical inhibitory circuit function and the development of stimulation strategies for sensory neuroprosthetic applications.

## Results

To investigate how PV and SOM inhibitory neuron subtypes respond to microstimulation and influence network activity, we performed two-photon calcium imaging and electrical stimulation in the somatosensory cortex of PV-Cre (n = 4) and SOM-Cre (n = 4) mice (Fig. 1A). In the canonical circuit, PV neurons provide fast-feedforward inhibition onto excitatory neurons, while SOM neurons provide slower-feedback inhibition onto excitatory and PV neurons, although these relationships vary across cortical areas (Fig. 1B)^27,33,40^. We found that the observed microstimulation-evoked responses were often interpretable within this framework. Virus was injected into the somatosensory cortex for expression of tdTomato in inhibitory subtypes and GCaMP in all neurons (Fig. 1C). Using a previously established classification approach^17^, we separated PV from non-PV neurons and SOM from non-SOM neurons based on tdTomato expression (Fig. 1D). PV neurons comprised ∼11% of recorded neurons in PV-Cre animals, while SOM neurons comprised ∼10% in SOM-Cre animals. Cortical neurons are ∼20% inhibitory, with PV a large fraction, though proportions vary across cortical regions^27,33,40^. Interestingly, SOM neurons exhibited significantly larger soma sizes (mean=299 µm^2^) compared to non-SOM neurons (mean=236 µm^2^, *p <* 0.001), while PV neuron soma sizes were not significantly different from those of non-PV neurons (*p =* 0.22, Fig. 1E). The SOM population recorded here may consist largely of larger Martinotti-type neurons, previously characterized as providing feedback inhibition that facilitates and increases supralinearly with input^41–43^.

**Figure 1.**
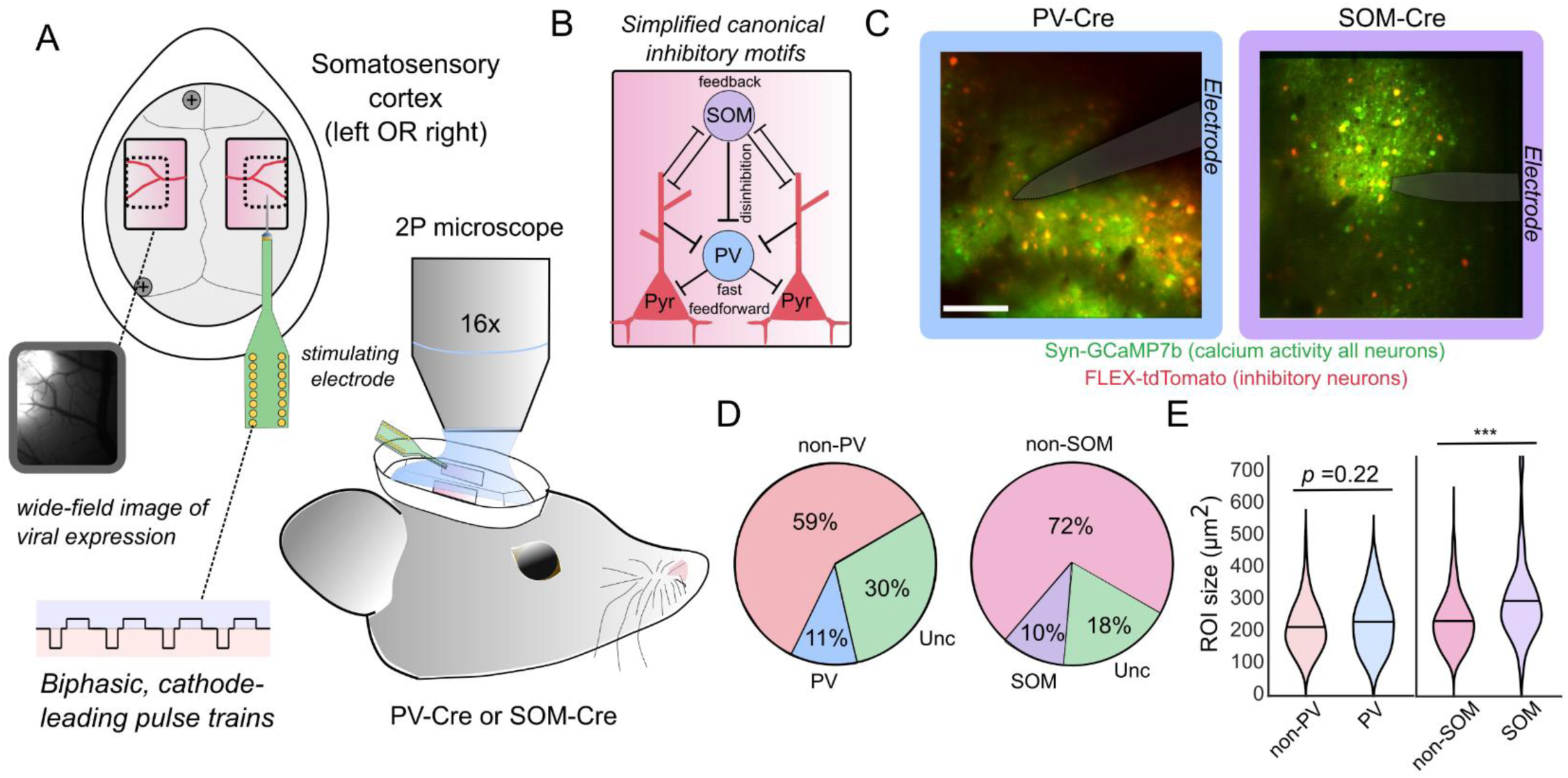
Two-photon imaging was conducted in the forelimb cortex of SOM-Cre and PV-Cre animals in response to microstimulation. (A) Craniotomies were performed bilaterally on SOM-Cre or PV-Cre animals. Stimulating electrodes were inserted into the forelimb sensory cortex based on anatomical landmarks and biphasic cathode-leading patterns were delivered and activity was imaged. The inset shows an image of virus expression in an injected mouse scanned with a wide-field microscope. (B) Canonical inhibitory motifs, with PV neurons providing fast feedforward inhibition onto pyramidal somata and SOM neurons providing feedback inhibition onto pyramidal dendrites and onto PV neurons. (C) Representative fields of view for PV-Cre and SOM-Cre animals, with Syn-GCaMP7b in green, FLEX-tdTomato in red, and the electrode shaded. Scale bar 100 µm. (D) Proportions of labeled, unlabeled, and unclassified (Unc) neurons in each animal model based on separation with linear regression. (E) Soma cross-sectional area, larger in SOM than non-SOM neurons and indistinguishable between PV and non-PV neurons. Group comparisons used linear mixed-effects models with neuronal subtype as a fixed effect and animal as a random effect. Model significance was assessed using analysis-of-variance tests of neuronal subtypes, and asterisks denote *p* < 0.05 (*), *p* < 0.01 (**), and *p* < 0.001 (***).

### Dynamic interplay between amplitude, frequency, and duration in shaping neural responses

Inhibitory neurons are more strongly activated by intracortical microstimulation^17^. High-frequency stimulation (100 Hz) also elicits strong responses in the first few seconds while producing weaker overall activity than low-frequency stimulation (10 Hz) across 30 s^13,16^. In humans, microstimulation frequency drives electrode-specific responses^5^, potentially reflecting distinct circuit recruitment. These findings prompted us to investigate how specific inhibitory neuron subtypes (PV and SOM interneurons) respond differently across a range of microstimulation frequencies. Frequency responses also depend on amplitude^5^. Therefore, each pattern was delivered at two amplitudes (10 and 20 µA). Our goal was to gain insight into how subtype-specific activation patterns shape network activity.

As expected, higher frequencies of microstimulation evoked strong neural activation within the first few seconds of microstimulation, followed by gradual reduction (Fig. 2A). Even 20-Hz stimulation (typically considered low frequency) elicited some response attenuation across 30 s. Only the 10-Hz condition produced a sustained and gradually increasing response. Consistent with prior findings, higher amplitude drove stronger activation than the lower amplitude (20 µA solid, 10 µA, dotted lines; Fig. 2A). The 100-Hz and 200-Hz response profiles were nearly identical. While calcium latencies limit resolution of small differences, the two patterns evoked similar responses despite 200 Hz delivering twice as many pulses. This pattern also aligns with previous perceptual reports, in which differences in lower frequency regimes (20 vs. 100 Hz) produce much more salient differences than higher frequency regimes (100 vs 300 Hz)^5^. At frequencies above 100 Hz, interpulse intervals approach the average membrane time constant (10 ms), potentially limiting further temporal integration by most neurons^12,44^.

**Figure 2:**
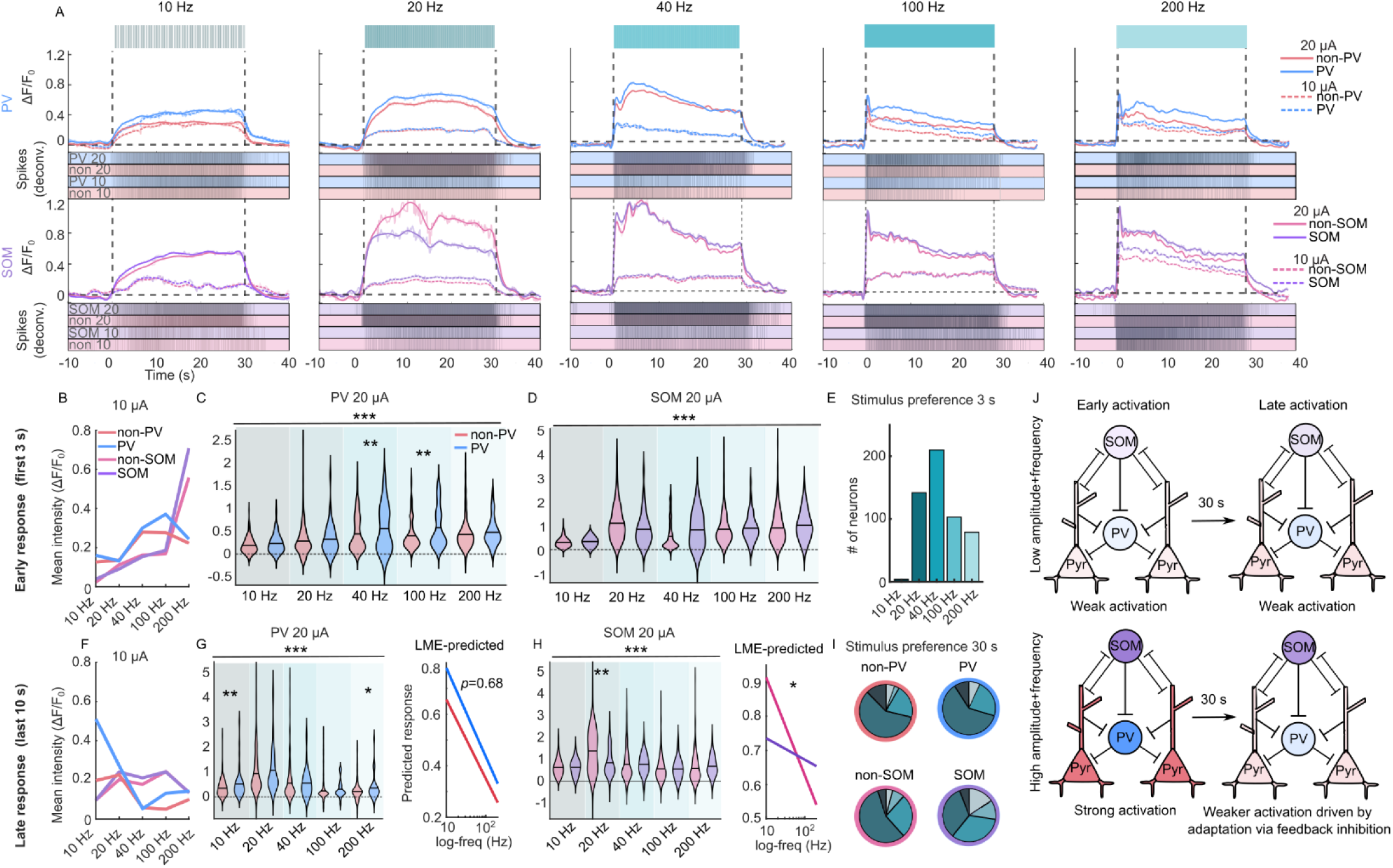
PV neurons generally respond stronger to microstimulation than non-PV neurons, with differences emerging with larger amplitude and duration, while SOM neurons are activated most strongly at 40 Hz and above. A) Representative traces of neural activity evoked by intracortical microstimulation at 10, 20, 40, 100, and 200 Hz, delivered at 10 and 20 µA. For each frequency, activity from PV (top) and SOM (bottom) animals is shown, with deconvolved spiking activity below. The solid line shows a spline fit overlaid on the underlying data in a lighter shade. (B-E) Mean fluorescence intensity during the first 3 s of stimulation. (B) Line plots showing mean responses at 10 µA for all cell types. Higher stimulation frequencies produce stronger activation at this low amplitude. (C) Violin plots comparing PV vs. non-PV neuron responses at 20 µA. PV neurons exhibit stronger activation, particularly at higher frequencies (*p* < 0.01 at 40 and 100 Hz). (D) Violin plots comparing SOM vs. non-SOM neurons at 20 µA. Different patterns evoked significantly different intensities of activation (*p* < 0.001) but none of the individual patterns evoked significantly different activation between SOM and non-SOM neurons (*p* > 0.05, all patterns). (E) Bar plot showing the distribution of frequency preferences (frequency producing maximal response) across all neurons. Most neurons respond strongest at 40 Hz, followed by 20 Hz, indicating that despite initially stronger activation at 100-200 Hz, rapid adaptation shifts overall preference toward intermediate frequencies within even short periods of time. (F-I) Mean fluorescence intensity during the final 10 s of stimulation. (F) Line plots showing 10 µA responses, where activity converges across frequencies due to adaptation and weak sustained activation. (G) Violin plots comparing PV vs. non-PV neurons at 20 µA, revealing greater sustained activation in PV neurons (*p <* 0.05 at 10 and 200 Hz). The right plot shows the LME-predicted values across log-transformed frequencies. The difference in PV and non-PV neurons does not vary by frequency (*p =* 0.68). (H) Violin plots comparing SOM vs. non-SOM neurons. Non-SOM neurons respond more at 20 Hz (*p <* 0.01). The right plot shows the LME-predicted values across log-transformed frequencies. There is a significant change in SOM vs. non-SOM neurons across frequencies, with non-SOM neurons having higher intensities at lower frequencies and SOM having higher intensities at higher frequencies (*p <* 0.05). (I) Pie charts showing frequency preference distributions for PV, SOM, non-PV, and non-SOM neurons at 20 µA. While PV, non-PV, and non-SOM neurons show similar distributions, SOM neurons exhibit a shift toward higher-frequency preferences (40-200 Hz), indicating that higher amplitudes and frequencies may result in stronger SOM-driven feedback inhibition over time. (J) Schematic illustrating proposed circuit dynamics. At low amplitudes and frequencies (top), stimulation evokes weak initial and sustained activation across the network. At higher amplitudes and frequencies (bottom), strong initial activation is followed by adaptation, likely driven by feedback inhibition from SOM neurons, leading to reduced pyramidal cell activity over time. Inhibitory (PV and SOM) neurons also adapt, but maintain relatively stronger activation to prevent runaway excitatory activity. Group comparisons used linear mixed-effects models with neuronal subtype as a fixed effect and animal as a random effect. Model significance was assessed using analysis-of-variance tests of neuronal subtypes with Benjamini-Hochberg correction for multiple comparisons, and asterisks denote *p* < 0.05 (*), *p* < 0.01 (**), and *p* < 0.001 (***).

In previous reports, neuronal activation weakened from onset to later phases of stimulation, hypothesized to reflect gradual buildup of inhibitory feedback^13,16^. To test this hypothesis, we analyzed responses separately during the first 3 s and the final 10 s of stimulation. Therefore, we hypothesized that at lower amplitudes, higher frequencies should drive greater activation because at near-threshold amplitudes higher frequency improves detection in primates^5,11^. Increasing frequency raised activity across all subtypes at 10 µA (Fig. 2B). Furthermore, most neurons had the strongest intensity evoked at 200 Hz in the first 3 s (Supp. Fig. 1A). However, low amplitudes of microstimulation drove weak, brief activation that adapts. By the last 10 s, activity was similarly weak across frequencies (Fig. 2F). This finding helps explain previous psychophysical results showing that higher frequencies improve detection near threshold and at short durations^5,11^, but that this relationship may weaken at higher amplitudes and longer durations.

Over short periods, higher frequencies would be expected to drive the strongest activation. Instead, at 20 µA, 40 Hz produced the strongest onset activation, despite delivering less than half the pulses of 100 or 200 Hz (Fig. 2C-E, Supp. Fig. 1B). Even within this short 3-s window, 100 and 200 Hz already showed some response attenuation (Fig. 2A), consistent with neural depression occurring within 1 s of microstimulation^16^. At 10 µA, 100 and 200 Hz produced stronger responses during the first 3 s aligning with expectations (Fig. 2B, Supp. Fig. 1A). Together, at lower amplitudes, higher frequencies drive activation more effectively, while at higher amplitudes, higher frequencies drive rapid depression of neural activity. These findings show that stimulation amplitude, frequency, and duration interact to shape the neural response to microstimulation.

### PV neurons are generally more activated by microstimulation than non-PV neurons, while SOM neurons are activated more strongly at higher stimulation intensities

Because neuronal subtypes have distinct temporal firing dynamics^27,33–35^, we hypothesized that microstimulation temporal patterns would recruit these subtypes differently. To test this, mean response intensity was compared across subpopulations at each microstimulation frequency. Within the first 3 s of microstimulation, inhibitory neurons were generally more strongly activated consistent with previous results^17^, although this effect was only significant for PV vs. non-PV at 40 and 100 Hz (*p <* 0.01, Fig. 2C,D). In the first 3 s, SOM neurons almost exclusively preferred the highest frequency, while other neuronal subtypes showed more diversity in their preferences (Supp. Fig. 1A). These results could be explained by strong activation of the network recruiting stronger feedback inhibition, resulting in strong SOM drive. Earlier work indicated that activity in excitatory neurons results in supralinear increases in inhibition, particularly Martinotti type neurons^43^. With optogenetic stimulation, the strong recruitment of feedback inhibition can result in paradoxical decreases in neural activity with excitatory neuron recruitment^45,46^. Together, these results motivated us to look into how the activity changes with continuous stimulation over time.

In the last 10 s of microstimulation, PV neurons had a higher mean intensity than non-PV neurons for all stimulation patterns (significant for 10 and 200 Hz *p <* 0.05). This stronger activation showed no significant changes across frequencies (*p =* 0.68, Fig. 2G, right). At 20 Hz and 20 µA, SOM neurons were significantly less activated than non-SOM neurons (Fig. 2H, *p <* 0.01). However, at higher frequencies, this trend reversed where SOM neurons were more strongly activated than non-SOM neurons (*p <* 0.05, Fig. 2H, right). This result suggests that high amplitude combined with high frequency preferentially recruits SOM neurons, while PV neurons are strongly activated for all stimulation input.

Additionally, 52-70% of PV, non-PV, and non-SOM neurons had the strongest mean activation at 20 Hz, while 67% of SOM neurons peaked at 40, 100, or 200 Hz (Fig. 2I). SOM neurons preferring higher frequencies tended to sit within 175 µm of the electrode, where activation is strongest (Supp. Fig. 1C). Collectively, these findings suggest different roles of PV and SOM neurons, where PV neurons provide strong feedforward inhibition to any input, while SOM neurons provide input-strength-dependent feedback inhibition that constrains excitatory activity, and that engagement depends jointly on frequency and amplitude.

These results highlight frequency and amplitude dependent interactions between inhibitory subtypes during microstimulation. Lower stimulation amplitudes and frequencies produced relatively weak and sparse network recruitment, which appeared insufficient to generate strong buildup of feedback inhibition over time (Fig. 2J). Under these conditions, neurons maintained activity throughout stimulation, preserving excitatory-inhibitory balance at lower activation intensity and likely lower perceptual salience. In contrast, higher intensities preferentially recruit SOM neurons, which may suppress the surrounding populations over time.

### 20-Hz stimulation shifts facilitated SOM neurons farther from the electrode and raises activation intensity with distance in non-inhibitory neurons

To better understand how neurons respond to fixed-frequency patterns, neurons were classified by how their activity changed over time, as previously described^17^. Classes included rapidly decreasing (RD), slowly decreasing (SD), no change (ND), increasing (Inc), and depressed (Dep). Neurons with increasing profiles were found previously to sit closer to the electrode, potentially reflecting sustained direct activation or calcium saturation, while distant neurons more commonly depressed, possibly reflecting synaptic fatigue or network-mediated suppression^14,17^. We therefore examined how the spatial distribution of these temporal response classes varied across inhibitory subtypes and stimulation conditions.

Consistent with prior observations, neurons exhibiting increasing activity profiles were more commonly found near the electrode (Fig. 3A,B, Supp. Fig. 1D). The relationship maintained across nearly all stimulation frequencies except 20 Hz at 20 µA. For this pattern, non-PV, non-SOM, and SOM neurons classified as increasing were located farther from the electrode than neurons classified as non-depressing, significant for SOM neurons (*p <* 0.01, Fig. 3C,D). In contrast, PV neurons with increasing activity profiles remained closer to the electrode, consistent with trends observed at other stimulation frequencies (Fig. 3C, Supp. Fig. 1D, *p <* 0.01).

**Fig. 3:**
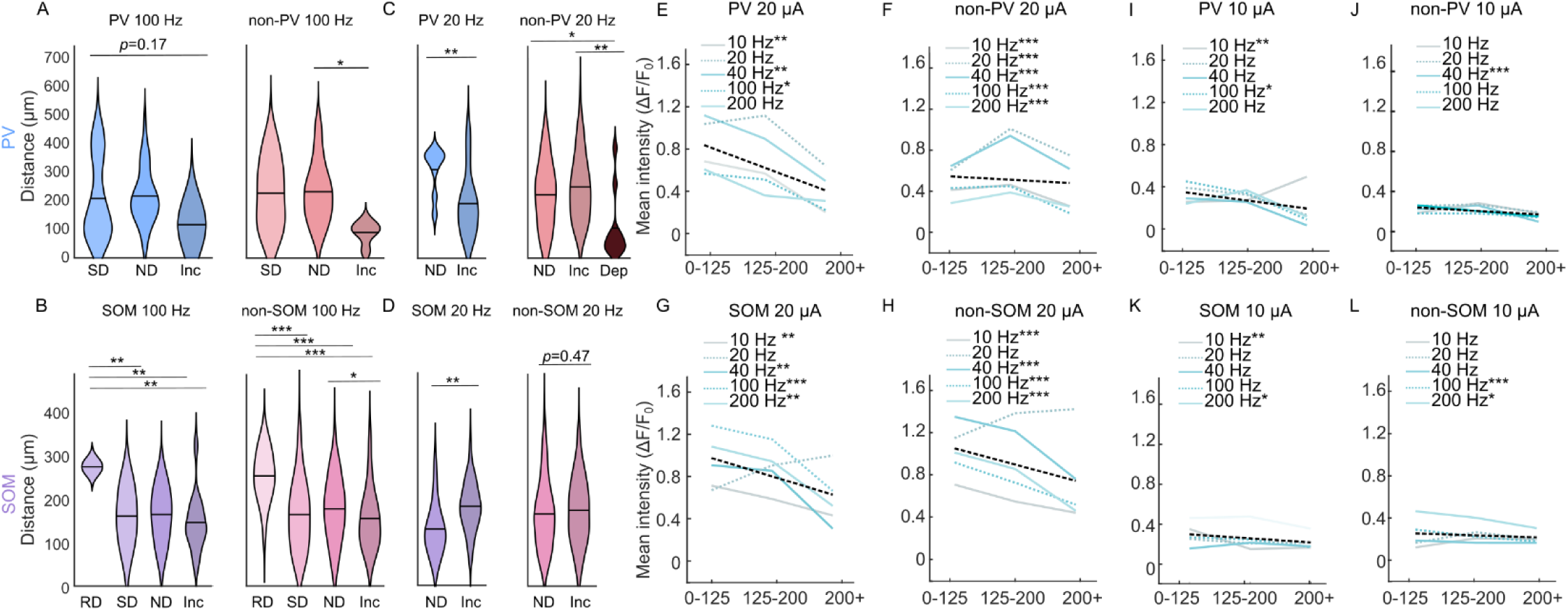
Stimulation frequency shifts the spatial profile of facilitation, while amplitude drives the distance-dependent falloff in neuronal activation. (A-D) Violin plots showing average distances of different neuron classes from the electrode. Typically, “increasing” neurons are closer to the electrode while “decreasing” neurons are farther from the electrode. For 20 Hz, the increasing neurons are farther from the electrode than the stable neurons for SOM (*p <* 0.01) and non-inhibitory neurons (*p >* 0.05), indicating more facilitation farther from the electrode. (E-L) Line plots showing the average fluorescence in three distance bins. The dotted line shows the average decrease across all plotted patterns. At 20 µA, intensity decreases with distance from the electrode, while at 10 µA it changes little across distance. SOM intensity decreased significantly across distance at every 20 µA frequency except 20 Hz. 20 Hz showed a non-significant increase across bins for SOM and non-SOM and a significant increase across the first two bins for non-PV neurons (*p <* 0.001). Group comparisons used linear mixed-effects models with either neuron class or distance bin as a fixed effect and animal as a random effect. Model significance was assessed using analysis-of-variance tests of neuronal subtypes with Benjamini-Hochberg correction for multiple comparisons, and asterisks denote *p* < 0.05 (*), *p* < 0.01 (**), and *p* < 0.001 (***).

These findings suggest that 20 Hz promotes sustained or facilitated recruitment at distal locations, except in PV neurons. One possible explanation is that 20 Hz represents an intermediate regime generating sufficient synaptic drive to facilitate network recruitment over time while avoiding strong synaptic depression or inhibitory buildup of higher frequencies. In contrast, lower frequencies such as 10 Hz may provide too little drive to sustain these effects. Lower amplitudes may do the same, since 20 Hz at 10 µA did not produce this effect (Supp. Fig. 1D). Interestingly, human perceptual studies report inconsistent effects at 20 Hz, where some electrodes evoke strong tactile sensations described as “tapping” and others evoke no detectable sensations at all^5^. The 20-Hz pattern may therefore depend on strong projections, providing strong synaptic drive with relatively weak direct activation of neurons.

### Stimulation at higher but not lower amplitudes drives stronger activation near the electrode

We next examined the activation intensity across three spatial distance bins (Fig. 3E-L). At 20 µA stimulation amplitudes, activation intensity was generally highest near the electrode and decreased in the most distal bin (Fig 3E-H). Neurons farther from the electrode are less likely to be strongly activated, which matches classical activation-distance relationships^6^. However, 20-Hz stimulation again produced a distinct pattern in SOM, non-SOM, and non-PV neurons, with activation intensity tending to increase at farther distances from the electrode, significant in the non-PV populations (*p <* 0.001, Fig. 3F).

In contrast, at 10 µA, activation intensity changed far less across distance bins while remaining above baseline in every bin, though some conditions still declined (Fig. 3I-L). This distributed recruitment differs from classical activation-distance models yet aligns with the sparse, spatially distributed recruitment described at low stimulation amplitudes^21,47^. These findings suggest that higher stimulation amplitudes produce dense local recruitment, while lower amplitudes drive sparse distributed activation through nearby low-threshold axons projecting to distal neurons.

These observations reconcile conflicting reports of spatial recruitment and separate the low- and high-amplitude stimulation regimes physiologically. In particular, lower-amplitude stimulation may produce more naturalistic network recruitment by avoiding the dense, synchronized local activation typical of microstimulation. Such stimulation strategies may also reduce the buildup of inhibitory activity and network depression that can emerge during prolonged or high-intensity stimulation.

### Burst patterns drive stronger activity over time than fixed frequency patterns

Previous studies have demonstrated that burst stimulation paradigms can be more effective than continuous, fixed-frequency microstimulation for sustaining perceptual responses over time^48^. In addition, different burst protocols are known to entrain neural populations to specific oscillatory rhythms^49^. Building on these results, our prior work indicated that theta-burst stimulation engages inhibitory neurons over extended periods^17^. Here, we examined how PV and SOM respond to different burst patterns (Fig. 4A).

**Fig. 4.**
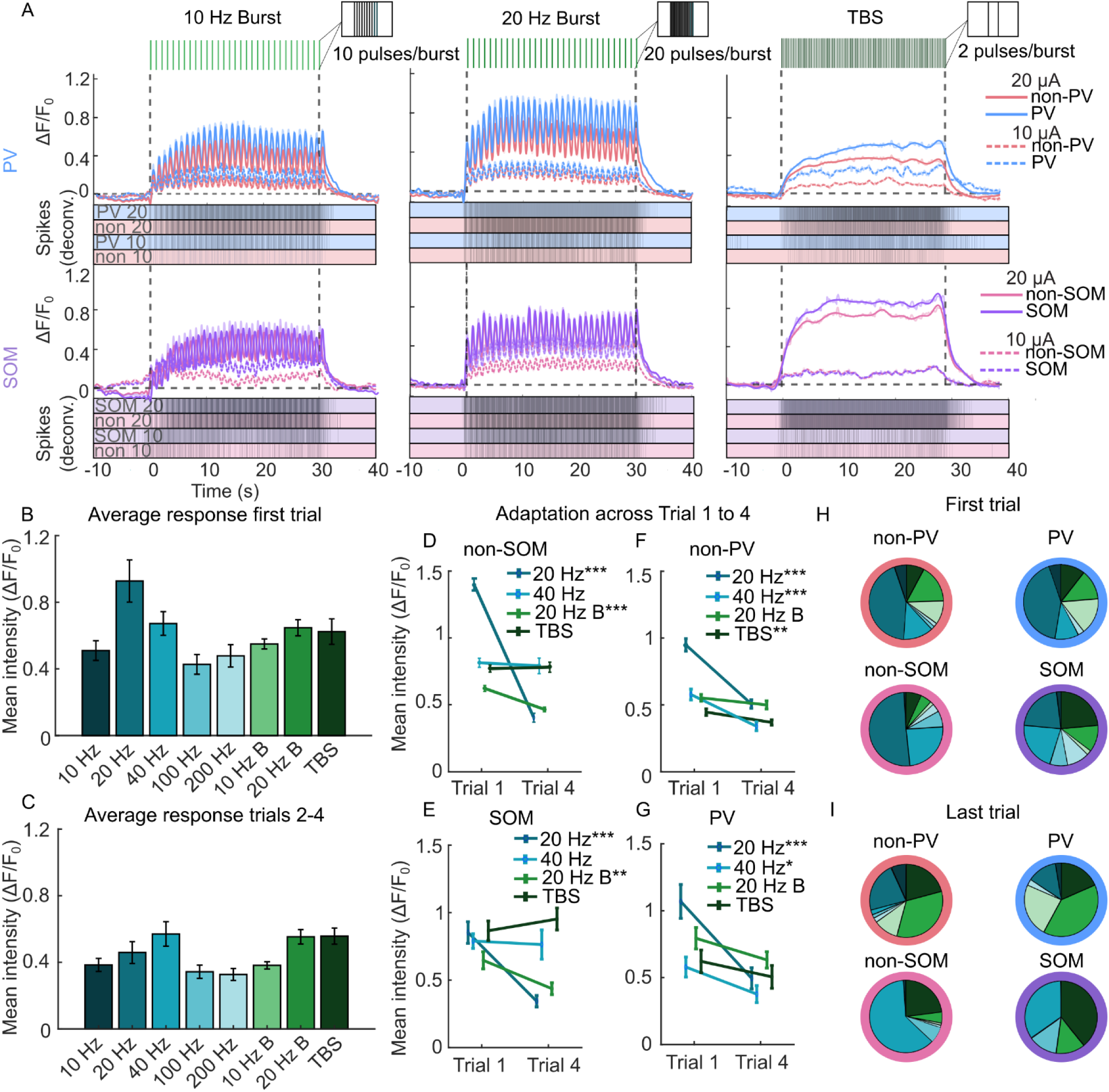
Burst-pattern stimulation maintains stronger activation across repeated trials than continuous high-frequency stimulation and TBS drives selective facilitation of SOM neurons. (A) Representative two-photon calcium traces showing neural responses to three burst stimulation patterns delivered at 10 µA and 20 µA: 10-Hz-Burst (10 Hz B), 20-Hz-Burst (20 Hz B), and theta-burst stimulation (TBS). For each pattern, activity is shown for PV (top) and SOM (bottom) neurons, with deconvolved spike rates below. The solid line shows a spline fit overlaid on the underlying data in a lighter shade. (B) Comparison of mean fluorescence intensity during the last 10 s of stimulation across all neurons for both frequency (from Fig. 2) and burst stimulation paradigms during trial 1. Continuous 20-Hz stimulation produced the strongest overall activation, while 40 Hz and the burst patterns elicited comparably strong sustained responses. (C) Comparison of mean fluorescence intensity during the last 10 s for trials 2-4. Here, 40 Hz, 20 Hz B, and TBS evoked the strongest responses, suggesting that burst patterns better maintain activation across repeated trials than continuous stimulation alone. (D-G) Mean activation from trials 1 and 4 for 20 Hz, 40 Hz, 20 Hz B, and TBS, separated by cell type (PV, non-PV, SOM, non-SOM). Continuous 20-Hz stimulation showed the strongest adaptation across trials, while 40 Hz and 20 Hz B exhibited smaller decreases. TBS showed minimal adaptation and even facilitation in SOM neurons (not significant) indicating TBS may be particularly effective at recruitment over time. (H-I) Pie charts showing the preferences of neurons to different stimulation patterns in the first trial (H) and the last trial (I). Within the first trial most neurons preferred the 20-Hz pattern, but 20 Hz caused the strongest adaptation across trials and results in weaker activation in later trials. SOM neurons however had the strongest preferences for 40-Hz and TBS patterns. By the last trial, neurons tended to have the strongest response to the 40 Hz or one of the burst patterns. PV neurons had a large majority of neurons that preferred burst-type stimulation (73.7%) while SOM neurons had a large representation of neurons that specifically preferred the TBS pattern (47.8%) in the last trial. Group comparisons used linear mixed-effects models with trial number as a fixed effect and animal as a random effect. Model significance was assessed using analysis-of-variance tests of neuronal subtypes with Benjamini-Hochberg correction for multiple comparisons, and asterisks denote *p* < 0.05 (*), *p* < 0.01 (**), and *p* < 0.001 (***).

Compared to fixed-frequency patterns, burst stimulation drove stronger late activity than high-frequency patterns, was comparable to 40 Hz, and was less effective than 20 Hz in the first trial (Fig. 4B). Neural activity decreased across trials, and in later trials, 20 Hz Burst and TBS patterns produced stronger responses than most patterns and were comparable to 40 Hz (Fig. 4C).

### SOM neurons are preferentially activated and facilitated by TBS across trials

While high-frequency fixed microstimulation patterns drive depression of neural activity and perception, previous work showed interleaved rest periods can mitigate this effect^16,17^. Therefore, we next looked at the adaptation to burst- and fixed-frequency patterns within and across trials. The 20-Hz, 40-Hz, TBS, and 20-Hz-Burst patterns were compared across trials, since these patterns produced the strongest overall activation (Fig. 4D-G). Interestingly, 20-Hz pattern activation adapted most from trial 1 to 4, despite driving sustained activation and distal facilitation in the first trial (*p <* 0.001). Activation with TBS, 20 Hz B, and 40 Hz was more stable from trial 1 to 4. TBS also increased activation from trial 1 to 4 in SOM neurons, although this effect was not significant (*p* > 0.05, Fig. 4E). This divergence suggests two distinct timescales and mechanisms, in which 20 Hz maintains activation within a pattern while depressing across trials, and TBS maintains or facilitates activation within and across trials, particularly for SOM neurons.

Activation intensity decreased for all other 20-µA patterns from trial 1 to 4 (Supp. Fig. 2A), while 10-µA pattern activation intensity remained stable or increased (Supp. Fig. 2B), supporting greater adaptation under strong stimulation. Neuron “preferences” for both fixed-frequency and burst patterns were evaluated across populations. Initially, most neurons preferred 20 Hz, while SOM neurons showed a stronger preference for TBS (27.5% vs 7.7% in non-SOM neurons, Fig. 4H). By the final trial, burst patterns and 40 Hz dominated preference. PV neurons favored burst patterns (73.7%), particularly 20-Hz bursts (34.2%), while SOM neurons showed a stronger preference for TBS (47.8% vs 23% in non-SOM neurons, Fig 4I). These findings suggest that burst stimulation better sustains activity across trials, while still strongly driving PV neurons, with SOM neurons particularly responsive to TBS.

20-Hz stimulation promoted facilitation in more distal neurons (Fig. 3D). TBS produced similarly enhanced activation, raising the question of whether TBS also preferentially recruited facilitating neurons distally. Increasing SOM neurons were located farther from the electrode on average than non-decreasing SOM neurons under TBS at 10 and 20 µA, although this effect was not significant (*p* > 0.05, Supp. Fig. 2C). Non-SOM neurons showed no such relationship, and PV and non-PV neurons showed the opposite, with increasing neurons closer to the electrode (Supp. Fig. 2C). This result further supports that TBS selectively promotes sustained facilitation of distal SOM neurons, which points to transsynaptic mechanisms, although a lack of significance prevents us from drawing a definitive conclusion. Together, these findings highlight that stimulation frequency and temporal patterning regulate inhibitory circuit recruitment differently, and burst delivery separates the PV and SOM neurons in time as well as in space.

### PV neurons are more adapted by strong fixed input over time, but more facilitated by burst type stimulation than unlabeled neurons

We next looked at adaptation within the first trial of stimulation (Fig. 5A,B, Supp. Fig. 3A,D). To quantify this effect, we used a change index, which measured the difference in neuronal activity between the second half (15-30 s) and the first half (0-15 s) of the stimulation pattern. PV changes exceeded non-PV changes in both directions: if non-PV neurons were depressed, PV neurons were more depressed and if non-PV neurons were facilitated, PV neurons were more facilitated (Fig. 5A, Supp. Fig. 3A-C). SOM temporal profiles resembled non-SOM profiles except at 20 Hz with SOM neurons having less facilitation than non-SOM neurons (Fig. 5B). This difference reached nominal significance (*p* = 0.029) but did not survive correction for multiple comparisons (*p* = 0.23, Benjamini-Hochberg criterion). In the last trial, SOM neurons tended to adapt and facilitate more than non-SOM neurons (Supp. Fig. 3E,F), particularly for the burst patterns. These findings indicate that burst patterns may be particularly strong at driving activation and facilitation of PV neurons while limiting build-up of SOM mediated inhibition until later trials.

**Fig. 5:**
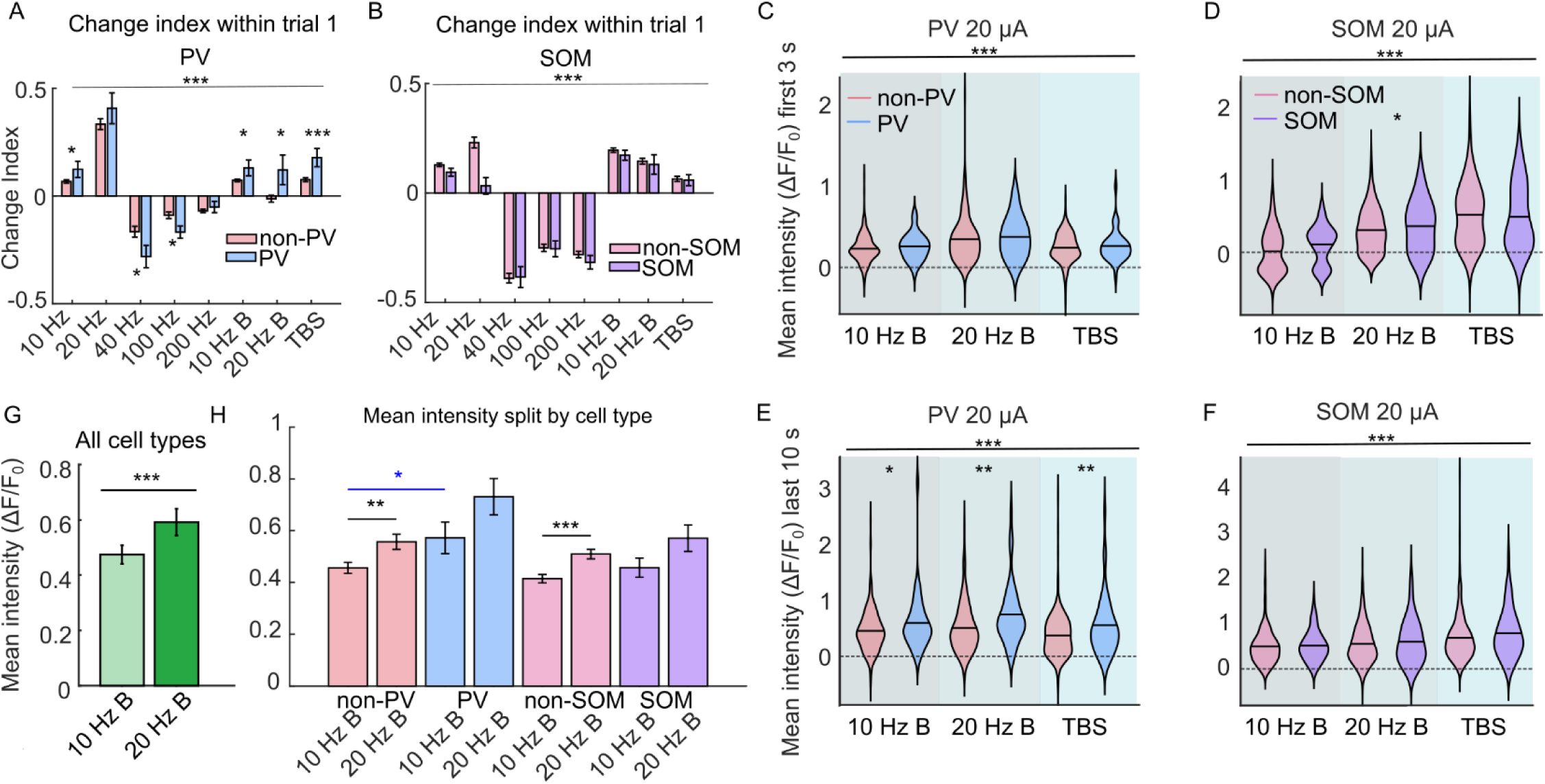
Burst patterns strongly activate PV neurons, with higher burst frequencies increasing activation across all subtypes. (A-B) Bar plots comparing PV vs. non-PV (A) and SOM vs. non-SOM (B) neurons across frequencies and burst patterns at 20 µA for the change index during trial 1. Lower frequencies (10 Hz, 20 Hz) and burst patterns generally showed facilitation (positive change index), while higher frequencies (40-200 Hz) exhibited adaptation (negative index). PV neurons demonstrated greater facilitation than non-PV neurons for the burst patterns (p < 0.05) and greater adaptation for high continuous frequencies (*p* < 0.01). SOM were not generally different than non-SOM neurons in the first trial, although differences were highly significant across different patterns (*p* < 0.001). (C-D) Violin plots showing mean fluorescence in the first 3 s for PV (C) and SOM (D) neurons under burst stimulation. The 20-Hz Burst produced significantly stronger activation in SOM than non-SOM neurons (*p <* 0.05). (E-F) Violin plots showing mean fluorescence during the final 10 s of stimulation for PV (E) and SOM (F) neurons. PV neurons exhibited stronger sustained activation than non-PV neurons for all burst patterns (*p* < 0.05), while SOM and non-SOM neurons showed no significant differences, although again difference were highly significant across patterns (*p* < 0.001). (G) Bar plot comparing all neurons’ responses to 10 Hz B and 20 Hz B, revealing significantly stronger activation with 20 Hz B (*p <* 0.001), indicating that higher burst frequencies can increase activation intensity without inducing additional adaptation. (H) The same comparison separated by cell type. Enhancement at 20 Hz B reached significance in non-PV and non-SOM neurons (*p* < 0.01). Group comparisons used linear mixed-effects models with either neuron subtype or stimulation pattern as a fixed effect and animal as a random effect. Model significance was assessed using analysis-of-variance tests of neuronal subtypes with Benjamini-Hochberg correction for multiple comparisons, and asterisks denote *p* < 0.05 (*), *p* < 0.01 (**), and *p* < 0.001 (***).

### Burst patterns strongly recruit PV neurons

Burst patterns changed the relative adaptation across subtypes, but to understand how this changed overall activation, fluorescent intensities were compared across subtypes for each burst pattern (Fig. 5C-F). In the first 3 s, PV and SOM neurons differed from unlabeled neurons only with the 20-Hz-Burst pattern and only in the SOM cohort (Fig. 5C,D), since bursting patterns require longer to build up subtype differences. However, bursting patterns differed in strength of activation, with 20 Hz Burst and TBS producing stronger activation than 10 Hz Burst (Fig. 5C,D, *p <* 0.001). TBS drove particularly strong activation in SOM neurons (Fig. 5D).

In the last 10 s, PV neurons were more strongly activated than non-PV neurons for 20-Hz-Burst and TBS stimulation (Fig. 5E, *p <* 0.05). Differences across patterns were still highly significant, with the strongest activation of PV neurons driven by 20-Hz-Burst stimulation (Fig. 5E, *p <* 0.001). None of the patterns produced differences between SOM and non-SOM neurons, although the differences across patterns were highly significant with TBS driving the strongest activation (Fig. 5F, *p <* 0.001). Consistent with the fixed frequency patterns, PV neurons were driven more strongly than non-PV neurons. Strong-amplitude, high-frequency bursts may drive strong PV activation without adaptation. SOM neurons did not have higher mean intensity than non-SOM neurons for any of the burst patterns in the first trial. This again supports that SOM neurons require strong continuous input that builds over time. Burst patterns, by introducing “duty cycles” in stimulation, may limit SOM-mediated feedback inhibition, unlike the continuous high-amplitude, high-frequency patterns (Fig. 2H).

### Higher burst frequencies enhance activation without increasing network depression

Beyond investigating how burst-patterns compared to fixed-frequency microstimulation, we also examined how individual bursts can be modulated to improve network activation. To determine whether increasing the frequency within individual bursts would enhance activation or instead exacerbate depression we compared burst patterns with 100 or 200 Hz intra-burst frequencies (Fig. 5G,H).

The 20-Hz Burst pattern (100-ms bursts of 20 pulses) produced stronger and more sustained network activation than the 10-Hz Burst pattern (100 ms bursts of 10 pulses) in every subtype (Fig. 5G, *p <* 0.001). With appropriate duty cycles, raising intra-burst frequency enhances network engagement without depression seen during continuous high-frequency stimulation. Burst-based microstimulation therefore has the potential to raise recruitment while preserving network stability.

### PV neurons reach peak activation faster during individual bursts, while SOM neurons peak more slowly than other cell types

In previous work, time-to-peak within bursts differed between inhibitory (VGAT+) and excitatory neurons^17^. Inhibitory neurons reached peak activation more rapidly near the electrode but more slowly at greater distances. We hypothesized that those spatial differences were attributed to distinct recruitment mechanisms with proximal neurons activated directly while distal neurons recruited through synaptic transmission. The delayed distal responses of inhibitory neurons indicated feedback inhibitory circuits.

The current findings suggest that SOM and PV interneurons account for these dynamics, consistent with their proposed roles in feedback and feedforward inhibition, respectively^27,28,32,33^. That framework predicts slower times to peak in SOM than non-SOM neurons at greater distances from the electrode, reflecting delayed recruitment through feedback pathways. Conversely, PV neurons should instead peak with or before excitatory neurons, consistent with rapid feedforward inhibition. Here, four burst paradigms (10 Hz Burst and 20 Hz Burst, each at 10 and 20 µA) produced clearly resolved single-burst responses against one in the previous study (10 Hz Burst 20 µA).

Consistent with this framework, SOM neuron responses reached peak activation more slowly than non-SOM neurons (Supp. Fig. 4B). This relationship was most clearly illustrated for the 20-Hz-Burst pattern at 20 µA (Fig. 6A). PV neurons peaked faster than non-PV neurons, although these relationships were noisier and the PV to non-PV difference was smaller than the SOM to non-SOM difference (Supp. Fig. 4A). Differences were largest in the 175 to 250 µm bins for PV neurons and in the 125 to 275 µm range for SOM neurons (Supp. Fig. 4).

**Fig. 6.**
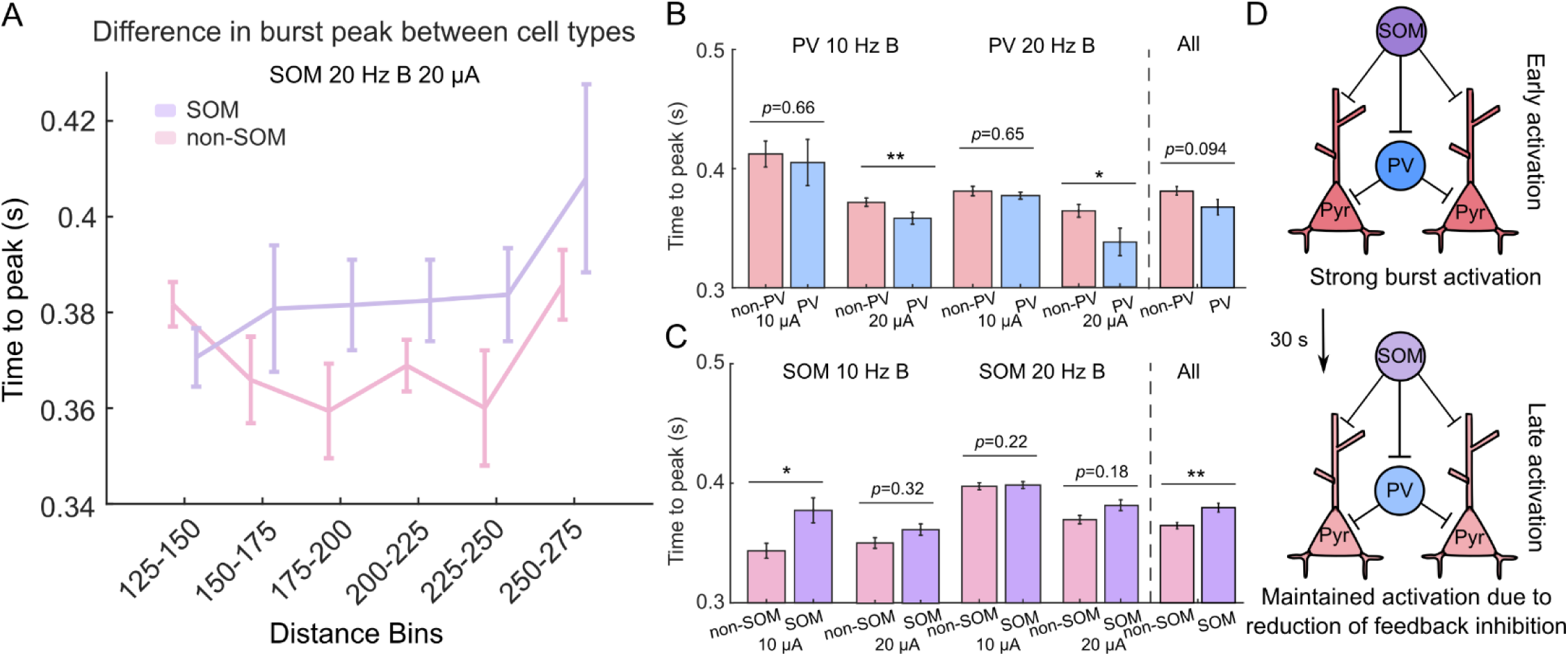
Distinct burst response timings of PV and SOM neurons support their proposed roles in feedforward and feedback inhibition, consistent with a model in which interburst intervals may limit the buildup of feedback inhibition. (A) The time it takes to reach the peak of the burst in the last six bursts for SOM vs. non-SOM neurons in response to the 20-Hz-Burst pattern at 20 µA. SOM neurons tend to have slower times to peak particularly in the farther distance bins. (B-C) Bar plots of (B) PV vs. non-PV and (C) SOM vs. non-SOM neurons farther from the electrode for each pattern as well as all patterns combined. PV neurons have faster times to peak than non-PV neurons and SOM neurons have slower times to peak than non-SOM neurons across all tested patterns, with two of the individual comparisons significant for PV and one for SOM (*p* < 0.05). Combining across all patterns revealed a significant difference between SOM and non-SOM (*p* < 0.01) but no significant effect for PV (*p* = 0.094). (D) Schematic illustration of network dynamics during burst-pattern microstimulation. Burst patterns evoke strong initial activation that is better maintained over time than continuous stimulation, likely due to reduced feedback inhibition during inter-burst intervals. Group comparisons used linear mixed-effects models with neuron subtype as a fixed effect and animal as a random effect. Model significance was assessed using analysis-of-variance tests of neuronal subtypes with Benjamini-Hochberg correction for multiple comparisons, and asterisks denote *p* < 0.05 (*), *p* < 0.01 (**), and *p* < 0.001 (***).

We next compared time-to-peak between PV and non-PV neurons, and between SOM and non-SOM neurons, in the spatial bins where the differences were largest. Across all burst paradigms, PV neurons consistently reached peak activation earlier than non-PV neurons, although this difference reached statistical significance only for the 10-Hz-Burst and 20-Hz-Burst pattern at 20 µA (Fig. 6B, *p <* 0.05). Conversely, SOM neurons consistently exhibited slower times to peak than non-SOM neurons, with statistical significance observed for the 10-Hz-Burst pattern at 10 µA (Fig. 6C, *p <* 0.05). For all cell types, 20 µA produced faster times to peak than 10 µA, potentially reflecting stronger synaptic recruitment and more reliable downstream activation. When combining across all patterns, the same overall trend appeared. However, only the comparison between SOM and non-SOM neurons reached statistical significance (*p* < 0.01), while the comparison between PV and non-PV neurons did not (*p* = 0.096). These results are consistent with the hypothesis that SOM neurons exhibit slower time-to-peak responses than non-SOM neurons, while PV neurons are faster or comparable to non-PV neurons. Altogether, these results support distinct temporal roles for these two interneuron populations in feedforward and feedback inhibition.

Together with our earlier findings^17^, this temporal separation may help explain how burst stimulation enhances neural activation both within and across trials relative to fixed-frequency patterns. Intermittent delivery of bursts may allow substantial network activation before delayed feedback inhibition develops. Overall, these results indicate that burst stimulation enhances network activation over time in part by limiting strong SOM-mediated feedback inhibition, despite strong facilitating activation (Fig. 6D).

### Dynamic amplitude modulation reduces build-up of inhibitory activity

Modulating microstimulation amplitude to mimic natural patterns of neural activation (described as *biomimetic stimulation)* can enhance perceptual quality and discriminability in humans^50,51^. This type of stimulation has also been shown to reduce synaptic depression and potentially improve the encoding of task-relevant information for brain-computer interfaces^16^. However, the impact of biomimetic amplitude modulation on inhibitory neuron activation has not been directly studied.

Dynamic amplitude modulation drove only brief transient activation rather than prolonged activation, potentially reducing inhibitory recruitment over time and preserving excitatory-inhibitory balance (Fig. 7A). Mean intensity did not differ between inhibitory and non-inhibitory subtypes (Fig. 7B, *p =* 0.81 for PV vs. non-PV and *p =* 0.70 for SOM vs non-SOM). Additionally, dynamic amplitude modulation reduced the adaptation of activity both within and across trials, agreeing with previous results (Fig. 7C-D). It also increased activity from trial 1 to 4 for PV neurons with BioAmp 100 Hz and for SOM neurons with BioAmp 200 Hz (*p <* 0.05, Fig. 7C), indicating that amplitude modulation can potentially improve excitability across trials.

**Figure 7:**
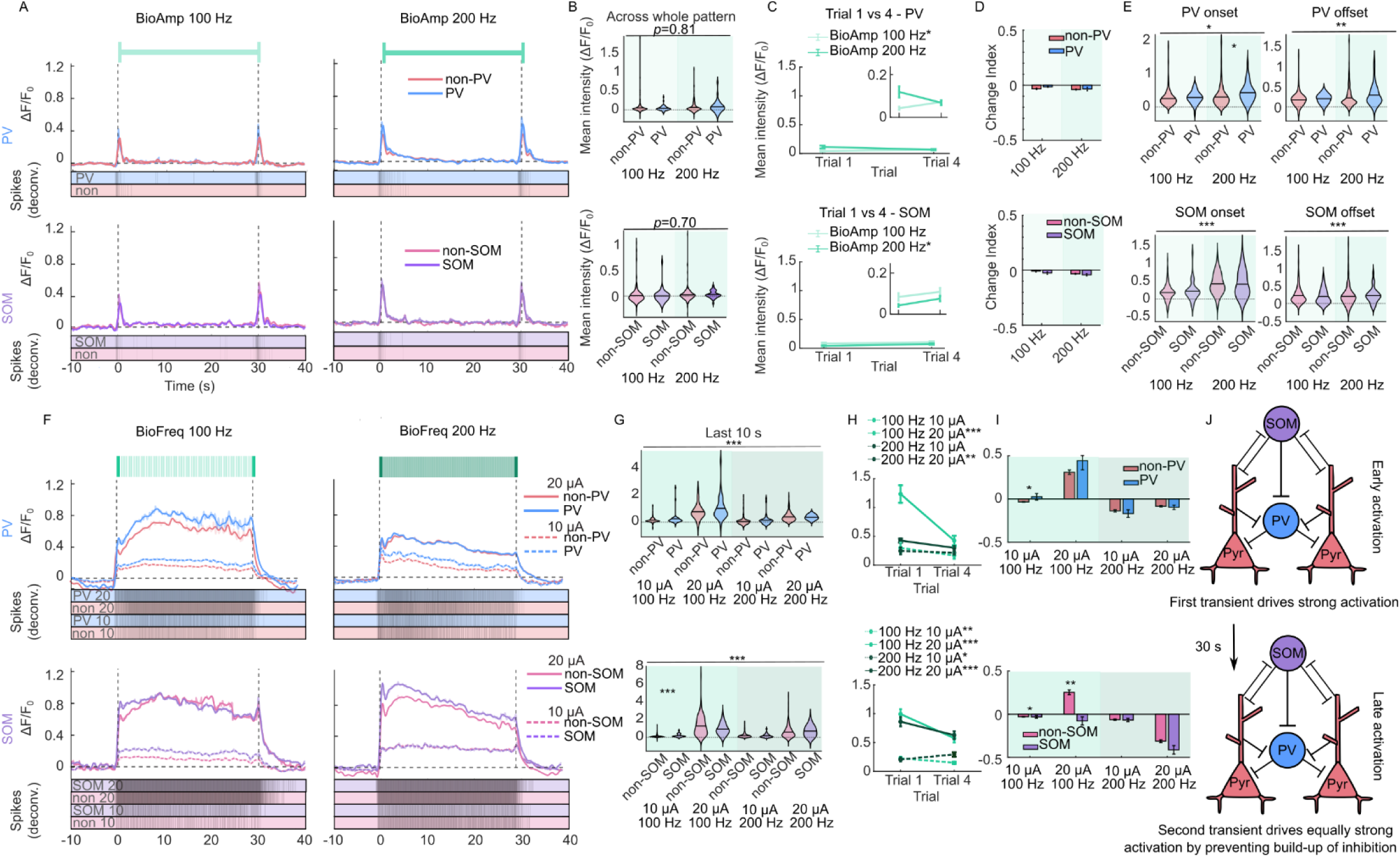
Dynamic amplitude modulation reduces adaptation and maintains excitatory-inhibitory balance, while dynamic frequency modulation evokes responses similar to fixed-frequency stimulation. (A) Representative two-photon calcium traces showing neuronal responses to two dynamic amplitude stimulation patterns delivered at 100 and 200 Hz. Both patterns include 100-ms onset and offset transients at 20 µA, with the remainder of the pattern delivered at 10 µA. For each pattern, activity from PV (top) and SOM (bottom) neurons is shown, with corresponding deconvolved spike rates below. The solid line shows a spline fit overlaid on the underlying data in a lighter shade. Strong activation is observed at both stimulus onset and offset. (B) Mean fluorescence across the full stimulus duration, averaged across all neurons, for both dynamic amplitude patterns. Neither pattern produced an imbalance between excitatory and inhibitory neuronal populations (*p* > 0.05). (C) Mean activation during trials 1 and 4, separated by cell type (PV, SOM). Dynamic amplitude modulation produces minimal adaptation across trials for all cell types, indicating preserved excitatory-inhibitory balance over repeated stimulation. The inset shows a zoomed in version to visualize the differences in pattern responses. For BioAmp 100 Hz in PV and BioAmp 200 Hz in SOM neurons, a small but significant increase in activation occurred across trials (*p* < 0.01), indicating sparse activation can potentially increase activation intensity over time. (D) Bar plots comparing PV versus non-PV neurons (top) and SOM versus non-SOM neurons (bottom) for the dynamic amplitude patterns, quantified using the change index during trial 1. Dynamic amplitude modulation induces little within-trial adaptation across cell types. (E) Violin plots of mean fluorescence during stimulus onset (first 3 s) and offset (last 3 s). During these transient periods PV neurons show significantly stronger activation than non-PV neurons (*p <* 0.05 at the 200-Hz transient frequency, while SOM and non-SOM neurons do not differ). There was a significant difference across patterns in all populations, with BioAmp 200 Hz generally producing stronger activation. (F) Representative two-photon calcium traces showing responses to four dynamic frequency stimulation patterns delivered at 10 and 20 µA. Each pattern includes a 100-ms transient at either 100 or 200 Hz followed by a hold period at 20 or 40 Hz respectively. PV (top) and SOM (bottom) activity is shown with corresponding deconvolved spike rates. The solid line shows a spline fit overlaid on the underlying data in a lighter shade. Weak offset transients are apparent for the 100-Hz patterns (left) but not for the 200-Hz patterns (right). (G) Violin plots of mean fluorescence during the final 10 s of stimulation for PV (top) and SOM (bottom) neurons. Similar to fixed-frequency stimulation, PV neurons show stronger activation than non-PV neurons, although no individual comparison were significant. Non-SOM neurons are more active than SOM neurons for the 100-Hz, 20-µA pattern with a 20-Hz hold period, consistent with Fig. 2H, although this difference was not significantly different after multiple comparisons correction. The 100-Hz, 10-µA pattern, however, produced significantly stronger activation in SOM neurons (*p* < 0.001). (H) Mean activation during trials 1 and 4 for PV and SOM neurons across all dynamic frequency patterns. Unlike dynamic amplitude modulation, dynamic frequency modulation does not reduce adaptation across trials. All 20-µA patterns produced significant adaptation from trial 1 to 4 (*p* < 0.01). (I) Bar plots comparing PV versus non-PV neurons (top) and SOM versus non-SOM neurons (bottom) for dynamic frequency patterns, quantified by the change index during trial 1. Increased adaptation is observed in SOM neurons for patterns dominated by a 20 -Hz hold period (*p* < 0.01), consistent with Fig. 5B. Together with (G), these results indicate that 20-Hz, 20-µA stimulation consistently evokes weaker SOM neuron activation across stimulation paradigms. (J) Schematic illustration of proposed network dynamics during dynamic amplitude modulation. Reduced stimulation amplitude during extended hold periods limits the buildup of inhibition, enabling strong network responses to both onset and offset transients. Group comparisons used linear mixed-effects models with neuronal subtype, trial number, or stimulation pattern as a fixed effect and animal as a random effect. Model significance was assessed using analysis-of-variance tests of neuronal subtypes with Benjamini-Hochberg correction for multiple comparisons, and asterisks denote *p* < 0.05 (*), *p* < 0.01 (**), and *p* < 0.001 (***).

Dynamic amplitude modulation drives the strongest activation in the brief transient periods, so we also compared the mean activation across subtypes for 1 s after onset and offset. The high-frequency (200 Hz) transients drove significantly stronger activation in PV neurons, agreeing with earlier results that higher frequencies and amplitudes quickly drive strong PV neuron activation (*p <* 0.05, Fig. 7E). Their brief duration, however, does not allow buildup of feedback inhibition, and SOM neurons were not significantly different than non-SOM neurons (Fig. 7E). Traditional microstimulation patterns may therefore induce atypically strong inhibitory activation, disrupting natural network dynamics. In contrast, dynamic amplitude modulation promotes a more physiologically relevant activation profile by lowering the buildup of inhibition in the network, potentially improving perception by maintaining more naturalistic circuit behavior.

### Low-Frequency dynamic frequency modulation enhances offset encoding but weakens onset encoding

In prior work, dynamic frequency modulation was less effective than amplitude modulation at encoding onset and offset transients^16^. Specifically, dynamic frequency modulation evoked strong adaptation of the network, resulting in an inability to drive strong offset transients. Those studies focused on higher-frequency regimes relevant to clinical use. However, lower-frequency modulation may improve offset encoding by reducing adaptation.

To test this hypothesis, two biomimetic frequency paradigms were compared: a high-frequency pattern (200-40-200 Hz) and a low-frequency pattern (100-20-100 Hz). As predicted, the lower-frequency modulation enhanced encoding of the offset phase, with more neurons showing offset responses compared to the high-frequency condition (Fig. 7F). However, dynamic frequency modulation did not match amplitude modulation in eliciting clear onset and offset responses. Additionally, in contrast to dynamic amplitude modulation, frequency modulation also failed to maintain excitatory-inhibitory balance. Similar to the fixed frequency profiles, inhibitory neurons were more strongly activated in most conditions, significant for SOM BioFreq 100 Hz 10 µA (*p* < 0.001, Fig. 7G). Activity evoked by 20-µA patterns also depressed strongly from trial 1 to 4 (*p <* 0.01, Fig. 7H), placing dynamic frequency modulation responses closer to fixed frequency stimulation than to dynamic amplitude modulation.

The BioFreq-100-Hz pattern primarily consisted of 20-Hz stimulation with brief 100-Hz onset and offset periods (Fig. 7F). Earlier, the pure 20-Hz pattern was the only fixed pattern that evoked stronger activity in non-SOM than SOM neurons (Fig. 2D,H). This pattern reproduced the stronger non-SOM than SOM activation, a difference that reached nominal significance (*p =* 0.043), but did not survive correction for multiple comparisons (*p =* 0.086, Benjamini-Hochberg criterion; Fig. 7G, bottom). At 20 µA, SOM neurons also showed significantly greater adaptation than non-SOM neurons under the BioFreq-100-Hz pattern (*p <* 0.001, Fig. 7I), matching the 20-Hz result. These consistent findings suggest that 20-Hz stimulation weakly recruits SOM neurons, reducing feedback inhibition and potentially sustain network activity.

Dynamic amplitude modulation, unlike frequency modulation, reduced adaptation within and across trials. Brief high-amplitude transients produced strong activation, while a low-amplitude hold preserved excitatory-inhibitory balance and sustained responsiveness. Although lowering frequency with dynamic frequency modulation reduced adaptation, it offered no advantage over continuously applying a fixed-lower frequency. Distance profiles support this principle, with dynamic amplitude (but not frequency) modulation producing much weaker changes across distance (Supp. Fig. 5). This distinction may help explain why dynamic amplitude modulation has shown greater promise for enhancing microstimulation perception and efficacy^52^. These findings support dynamic amplitude modulation as a promising strategy for improving microstimulation efficacy while limiting inhibitory buildup and preserving network activity.

### A simple computational model incorporating strong adaptive PV inhibition and time-dependent SOM inhibition reproduces responses across stimulation patterns

Throughout this paper, we placed our observations in the context of the canonical inhibitory circuit motif, in which PV neurons provide strong adapting inhibition onto the cell bodies while SOM provides facilitating inhibitory input onto the dendrites. We made a simple computational model using these principles to reproduce the observed responses (Fig. 8A). As observed throughout the paper, the activity in each subpopulation is similar in terms of shape for a given stimulation pattern. To reproduce this effect in our model, the key assumption was that the intrinsic properties of the individual neuron type does not produce distinct profiles nor does the strength of the input from excitatory neurons onto PV or SOM neurons change. Instead, the strength of PV and SOM input onto excitatory neurons changes over time (represented by F_SOM_ and D_PV_, Fig. 8A). This differential change in PV and SOM synapses is compatible with literature demonstrating that optogenetic-TBS induces long-term depression at PV neuron synapses onto pyramidal neurons via calcineurin-dependent pathways, while inducing long-term potentiation at SOM neuron synapses onto pyramidal neurons via CaMKII-dependent pathways^53^. It has also been shown in visual cortical slices that PV neurons can fire continuously over time while their input onto pyramidal cells adapts^54^.

**Figure 8:**
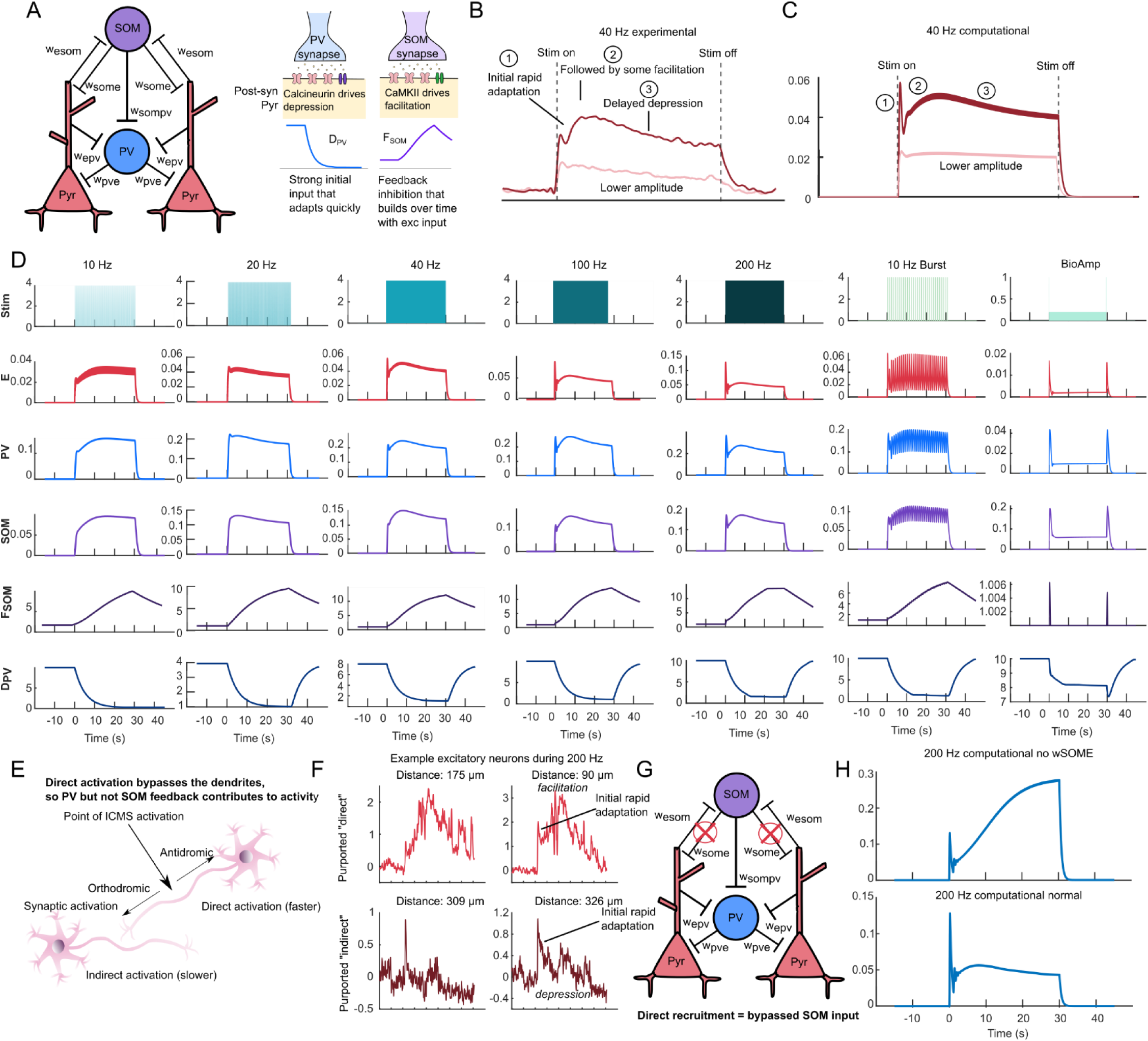
A simple recurrent rate-based network model reproduces the qualitative response dynamics across stimulation patterns. (A) A structural description of our model based on the canonical inhibitory motifs described throughout this paper. Critical to this model was the adaptation of PV synapses onto excitatory neurons and the facilitation of SOM synapses onto excitatory neurons. Previous work has shown that PV synapses can depress via calcineurin mediated effects while SOM synapses can facilitate via CaMKII effects^53^. (B) The average fluorescent trace of non-PV neurons responding to 40 Hz at 20 and 10 µA. The response to 40 Hz 20 µA had the most unique dynamic response, which is highlighted in this figure in three phases. We used this dynamic response as a reference in developing our model. (C) Our computational model reproduced the three stages of the dynamic response for 40 Hz microstimulation, and was also able to reproduce the effect of lower amplitude microstimulation in which the response is weaker and the unique dynamics disappear (since higher drive of the network is needed to drive these effects). (D) The computational model produced the characteristic shapes of the responses to the patterns used in this paper using the same weighting parameters across conditions. (E) A basic theoretical description of how microstimulation recruits neurons directly and indirectly. Neurons recruited directly are activated at the axon or axon hillock and propagate towards the cell body, which bypasses integration at the dendrites. (F) Example of two purported “direct” recruited neurons and “indirectly” recruited neurons. Both types of neurons experience initial adaptation, but directly recruited neurons are able to facilitate after initial adaptation while synaptically recruited neurons tend to depress over time at high amplitudes and frequencies. (G) From our model, directly recruited neurons should bypass SOM input which inputs onto the dendrites, while still receiving input from PV neurons that input onto the cell bodies. (H) Excluding SOM input in our model produces the expected “directly” recruited behavior, with initial adaptation but facilitating or “increasing” activity across the pattern.

The 40-Hz, 20-µA pattern produced the most complex dynamic profile, so we tested whether our model could reproduce this behavior (Fig. 8B). Activation began strongly, then showed rapid adaptation, some facilitation, and finally long-term depression. These complex dynamics disappeared at the lower amplitude, and the model reproduced this relationship (Fig. 8C). In the model, strong initial PV input drives rapid adaptation, which is then followed by facilitation once the PV input adapts. SOM synapses facilitate, then driving long-term depression. Therefore, this simple model of PV and SOM input informed by the literature reproduces the observed dynamic phenomenon, including the weaker and less dynamic response driven by lower amplitudes of microstimulation (Fig. 8C).

The model was expanded to incorporate the other tested frequencies, a burst pattern, and a dynamic-amplitude profile, and approximately reproduced the observed responses (Fig. 8D). Parameters were semi-optimized to reproduce experimental results, and the same weighting parameters were used for all patterns, meaning a single continuous model reproduced observed responses across patterns. Initially, the model only included direct stimulation of pyramidal neurons. When direct stimulation of PV and SOM neurons was added, the drive needed to optimally reproduce the results was 0.1-0.2, relative to 0.9 for excitatory neurons. This result suggests that inhibitory neurons are less strongly directly recruited by microstimulation than excitatory neurons under the conditions tested, with their responses arising predominantly through transsynaptic recruitment. One possible explanation is cell-type-specific differences in axonal morphology and excitability, which can influence the efficiency of extracellular stimulation.

### Differences in direct and indirect recruitment reflect bypassed synaptic integration and blocked SOM-mediated feedback inhibition, despite rapid PV-mediated adaptation at the cell body

In the present and previous work, neurons which “increase” across the stimulation pattern tend to be near the electrode, while neurons that “decrease” across the pattern tend to be farther from the electrode. This difference was previously attributed to direct versus indirect activation. For neurons that increase and are thought to be directly recruited (Fig. 8E), we still observe initial onset depression^17,21^. Example neurons at 200 Hz near the electrode showed initial adaptation followed by facilitation, interpreted as “directly” recruited (Fig. 8F). Neurons farther from the electrode showed the characteristic adaptation and depression, interpreted as “indirectly” recruited (Fig. 8F).

A recent review clarified that neurons recruited directly are recruited at the axon or axon hillock driving activation of the neuron via antidromic activation, which bypasses integration of input at the dendrites^12^ (Fig. 8E). SOM neurons synapse onto the dendrites, so they affect indirectly but not directly recruited neurons, while PV neurons synapse onto the cell body and affect both (Fig. 8G). Removing SOM input from the model reproduced these effects. Initial adaptation driven by PV input, followed by facilitation across the pattern (Fig. 8H). This mechanism can explain why increasing neurons tend to sit near the electrode (more likely to be directly recruited) while decreasing neurons are farther away (more likely to be synaptically recruited).

Altogether, this simple explanatory model reproduced many of the dynamic responses observed in our experiments and provided a mechanistic framework for interpreting them, including effects on directly vs. indirectly recruited neurons. The model was not intended as a predictive model, and we did not use it to prospectively test or validate the contribution of individual neuronal subtypes. Instead, it was constructed from known physiological roles based on prior literature and used to determine whether these mechanisms could account for the observed responses. This model further suggests that inhibitory neurons shape the response to microstimulation primarily through their synaptic effects, instead of through distinct intrinsic responses to stimulation, since the individual subtypes showed similar responses to microstimulation in our experiments.

## Discussion

We investigated how parvalbumin-positive (PV) and somatostatin-positive (SOM) inhibitory interneurons responded to intracortical microstimulation with distinct amplitude and frequency dependence across a broad range of temporal patterns. Our findings reveal that microstimulation recruits excitatory and inhibitory subnetworks in a frequency, amplitude, and duration dependent manner. Furthermore, PV and SOM interneurons contribute in unique and evolving ways to cortical activity during prolonged stimulation. Perceptual and physiological microstimulation effects therefore emerge from interactions between excitation, inhibition, disinhibition, and adaptation, rather than from direct activation alone.

### Low-amplitude, sparse recruitment preserves naturalistic dynamics while strong sustained drive accumulates inhibition

Stimulation frequency interacts strongly with amplitude and duration to shape network activation. At the lower amplitudes, increasing frequency generally increased neural activity during stimulation onset, consistent with the idea that more pulses more effectively recruit cortical networks when near threshold. This onset increase aligns with psychophysical studies showing higher-frequency microstimulation improves perceptual detectability at low amplitudes^5^. However, this relationship reversed at higher amplitudes and over longer durations. For the higher amplitude, intermediate frequencies (particularly 40 Hz) produced the strongest sustained activation, while 100 and 200 Hz rapidly depressed network activity after the onset (Fig. 2A,F). These findings suggest that strong activation at high frequencies quickly engages activity-dependent suppressive mechanisms. Strong PV activation in these early onset periods potentially indicates recruitment of feedforward inhibition from PV neurons that accompanies the observed rapid depression.

From multiple studies, an “optimal frequency” of microstimulation is unclear and depends on the preparation. High frequencies at high amplitudes drive depression over duration, so the combination of these three must be considered together as well as the desired outcome on the neural population. Here, for fixed frequency patterns, higher frequencies drove the strongest response in the first 3 s at 10 µA, while 40 Hz drove the strongest sustained response in the final 10 s at 20 µA (Fig. 2B,F). Other work that reported the strongest activation at low frequencies used 10-s patterns in anesthetized visual cortex at fixed 20 µA^55^, while the present recordings used 30-s patterns in awake forelimb cortex at two amplitudes. Cortical state can also influence which neurons are recruited by microstimulation^56^. Anesthetic state, cortical area, stimulus duration, and amplitude covary with the reported optimum, and no single study varied all of them.

In addition to temporal dynamics, the present findings provide insight into the spatial organization of microstimulation recruitment. At higher amplitudes, activation was strongest near the electrode and declined with distance (Fig. 3E-L), consistent with classical models of electrical stimulation^6^. By contrast, lower amplitudes produced relatively sparse and spatially distributed activation patterns with little distance dependence, closely resembling prior observations from low-amplitude microstimulation studies^21,47^. During these low-amplitude patterns, PV and SOM neurons were generally not significantly more activated than excitatory neurons, unlike the higher-amplitude patterns. These findings help reconcile conflicting reports of whether stimulation produces dense local recruitment or sparse distributed activation^6,47,57^. Computational modeling previously attributed sparse, distributed activation to axons projecting away from the electrode, with higher amplitudes densely activating the whole cortical column^57^. The present recordings confirm this physiological effect directly and show that the conflicting reports arise from different amplitude regimes. Sparse distributed recruitment may better preserve naturalistic cortical dynamics by avoiding excessive synchronization and inhibitory buildup, indicating that low-amplitude regimes used in moderation can benefit stimulation protocols.

### SOM recruitment depends on sustained drive rather than peak instantaneous rate

PV and SOM interneurons contribute differently to microstimulation temporal dynamics. During stimulation onset, PV neurons were generally strongly activated and showed pronounced adaptation at higher stimulation intensities during prolonged stimulation. In contrast, SOM neurons exhibited preferential recruitment during sustained high-amplitude and high-frequency stimulation, particularly at frequencies above 20 Hz. SOM neurons were less activated than non-SOM neurons at 20 Hz and 20 µA, but this pattern reversed at higher frequencies (Fig. 2H). By comparison, PV neurons exceeded non-PV neurons in the final 10 s for all frequencies (Fig. 2G). These findings support a model in which PV neurons contribute the fast inhibitory response to microstimulation, while SOM neurons become progressively engaged as network activity accumulates.

This interpretation is consistent with known circuit motifs in cortex. PV interneurons receive strong thalamocortical and local excitatory input and are well positioned to mediate rapid feedforward inhibition^27,28,33^. SOM interneurons, particularly Martinotti-type cells, are more strongly associated with recurrent cortical excitation and feedback inhibitory control^27,28,33^. Previous work has demonstrated that SOM recruitment can scale supralinearly with excitatory network activity and can strongly suppress pyramidal cell activity during sustained activation^43^. These findings suggest that prolonged or intense microstimulation preferentially drives this SOM-mediated feedback inhibitory regime, resulting in progressive suppression of cortical activity over time.

SOM somata in these recordings were larger than non-SOM somata (299 µm² against 236 µm², Fig. 1E), consistent with a population weighted toward Martinotti-type cells^41–43^. Supralinear recruitment predicts the intensity-dependent rise measured here (Fig. 2D,H). At 10 µA and 10 to 20 Hz, SOM neurons are only weakly activated, and their output does not visibly constrain the surrounding population. Above 20 Hz at 4 nC/phase, SOM activation rises and suppression of the surrounding population follows. Activation was weaker in SOM than in non-SOM neurons at 20 Hz within the first trial (Fig. 2H). The same weak drive reappeared in the BioFreq 100 Hz pattern, delivering 20 Hz between brief transients (Fig. 7G,I). Reduced SOM recruitment therefore depends on the sustained amplitude and frequency of a pattern rather than on its peak instantaneous rate.

### PV and SOM neuron activation separates in time and space

Response timing during burst stimulation supported the proposed feedforward role for PV neurons and the feedback role for SOM neurons (Fig. 6). Calcium imaging usually cannot resolve latency differences across cell types, but burst stimulation produced transient responses sharp enough for time-to-peak to serve as a proxy. Across burst paradigms, PV neurons consistently peaked earlier than non-PV neurons, while SOM neurons peaked later than non-SOM neurons, consistent with their proposed functional roles (Fig. 6C). Times to peak diverged beyond 175 µm from the electrode (Supp. Fig. 4), close to the 225-µm crossover reported previously for the VGAT-labeled population^17^. In these more distant bins, more neurons are expected to be activated by synaptic rather than direct activation.

An unexpected finding was that SOM neuron responses more closely resembled those previously observed in the broader VGAT-labeled inhibitory population^17^ than did PV neurons. The indicator, classification approach, response-class criteria, and awake preparation were matched to that study, narrowing the candidate explanations to cortical area and to subtype composition of the VGAT population. Greater involvement of SOM interneurons in layer 2/3 recurrent circuits of forelimb cortex may explain the difference. Consequently, the inhibitory response measured in the earlier VGAT response may not represent the weighted average of inhibitory subtypes but may instead be dominated by whichever interneuron population the paradigm engages most strongly. Comparing interneuron subtype recruitment across cortical regions and laminae will be important for determining the generality of these observations.

A simple computational model reproduced experimental results based on these dynamics, providing stronger evidence that the results can be interpreted in this framework. This model also provided three important insights. First, the effects of different temporal patterns are likely driven by changes at the synapse, which is why excitatory, PV, and SOM neurons show similar shapes of activation over time, despite each playing different roles in the propagation of stimulation evoked activity. PV neurons are known to have higher firing frequencies, and recent work showed they may be more responsive to high-frequency external electric fields over short periods. In that work, pyramidal neurons entrained most strongly near 8 Hz and PV neurons near 140 Hz, matching their intrinsic firing rates^35,58^. All neuronal subtypes had similar response profiles over time to different patterns, indicating that extended durations of ICMS result in more complex dynamics in which the synaptic effects become more important than the individual neuron response dynamics. Second, the model predicted weaker activation of PV and SOM neurons by microstimulation than pyramidal neurons, so that inhibitory effects are primarily mediated by synaptic dynamics. Third, the distance relationships, particularly ones in which neurons near the electrode increase in activity while distal neurons decrease in activity over time, can be explained from a basic recruitment standpoint, where stimulation bypasses dendritic input and therefore prevents SOM feedback from dampening responses in directly recruited neurons. Interestingly, the model depended on the output of PV neurons adapting over time and the output of SOM facilitating. The literature has more often shown these effects occurring at a different part of the pathway: synapses onto PV adapt and synapses onto SOM facilitate^41,43^. However, other more recent work supports that PV synapses onto pyramidal neurons can be adaptive while PV neurons fire consistently, and SOM synapses onto pyramidal neurons can be facilitating^53,54^. These findings together provide a strong explanatory framework for measured stimulation evoked responses, but further studies with additional targeted manipulations will be needed to validate this model.

### 20-Hz microstimulation window engages weak SOM recruitment

Interestingly, the present results identify 20 Hz as the frequency window that most effectively sustains network activation within the first trial, and two-photon recordings in anesthetized visual cortex showed the strongest sustained, activation-dominant response at 25 Hz^55^. Chronic recordings in awake visual cortex showed progressively larger Ca^++^ responses at 25 Hz growing across 12 weeks, and epileptiform events in 4 of 9 mice almost exclusively at 25 Hz, reaching ∼15% of trials by day 42 before plateauing. Here, 20 Hz in awake forelimb cortex sustained activation, facilitated distal neurons (Fig. 3D), and recruited non-SOM neurons significantly more strongly than SOM neurons (Fig. 2H). Consistent with this weak low-frequency SOM engagement, single-cell recordings showed that SOM neurons have instantaneous firing rates at intermediate to high frequencies (>30 Hz) with much lower rates of instantaneous firing at 0-20 Hz, correlating with their preferential response to intermediate frequencies of weak electrical fields^35,58^. Drive at 20 to 25 Hz recruits recurrent excitation while SOM dendritic feedback stays weak, which propagates activity within a pattern and leaves the network less stabilized across repeated sessions, which could drive epileptiform activity.

Electrical brain stimulation can be used therapeutically to reduce seizure burden in epilepsy^59,60^. It has been posited that stimulation can alleviate seizures by restoring excitatory-inhibitory balance, although inhibitory drive may also increase seizures depending on the context^59,61^. Therapeutic stimulation paradigms have commonly employed either low frequencies (<10 Hz) or high frequencies (>50 Hz), although the optimal frequency depends on the stimulation target and paradigm. Interestingly, 25 Hz was more likely to produce post-stimulation discharges with epileptiform features with microstimulation of visual cortex^55^. In other work, low-frequency electrical stimulation was shown to generate seizure-like activity, while this was less likely with high-frequency electrical stimulation, and this observation was posited to arise from high-frequency electrical stimulation recruiting PV neurons^62^. These findings raise the possibility that low or intermediate stimulation frequencies engage network dynamics permissive to prolonged or recurrent activity, by reducing the network inhibition that normally prevents runaway activation. This vulnerability was measured acutely, yet it connects to two chronic observations. The implanted electrode alone alters the excitability of nearby neurons over weeks, and chronic microstimulation can produce seizure-like activity^55,63^. A reduction in overall network excitability need not be protective. If chronic changes fall more heavily on PV and SOM neurons than on excitatory neurons, inhibitory restraint would decline relative to excitation^64,65^, the same imbalance that the weak inhibitory recruitment at 20 Hz produces acutely. Whether the chronic excitability changes are biased toward inhibitory neurons, and whether this bias raises seizure susceptibility at specific frequencies, remains untested. Future work combining chronic recordings with subtype-specific labeling across these frequencies could address this gap.

However, 20-Hz stimulation decreased activation across trials (Fig. 4D). An additional plasticity-dependent mechanism may make 20 Hz fail across many bouts^55^. Microstimulation was recently shown to produce plastic changes in the synapse over chronic time periods correlating with learning, indicating microstimulation can indeed engage plastic mechanisms, although it is unclear what role inhibition plays in these dynamics^66^. Unusual properties of 20-Hz stimulation may help explain the highly variable perceptual effects reported in prior human microstimulation studies at this frequency^5^. Some electrodes evoke clear tactile percepts described as tapping or flutter, while others fail to evoke detectable sensations altogether. These results suggest that 20-Hz stimulation may depend particularly strongly on the underlying circuit architecture and connectivity surrounding the electrode. Because 20 Hz recruits little direct inhibitory stabilization and relies on synaptic propagation, its effect depends on the projections and local subnetworks surrounding the electrode, which may contribute to the variable perceptual reports at this frequency in human microstimulation.

### Duty-cycling and dynamic-amplitude modulation sustain activation by limiting inhibitory accumulation

Burst-pattern stimulation substantially alters inhibitory recruitment and network adaptation. Burst paradigms maintained stronger activity over time than continuous high-frequency stimulation and reduced adaptation across repeated trials (Fig. 4B,C). These effects likely arise because inter-burst intervals provide opportunities for partial recovery from synaptic depression and inhibitory accumulation. Increasing the pulse count within a burst raised activation intensity without driving adaptation (Fig. 5E,F). The charge ceiling per burst remains unclear, so optimal burst intensity and duration are open, and amplitude modulation offers a different way to limit inhibitory buildup.

Perhaps the most translationally relevant finding for sensory restoration is that dynamic amplitude modulation was distinctively effective at preserving excitatory-inhibitory balance and minimizing adaptation (Fig. 7A,B). We previously showed that dynamic amplitude modulation but not frequency modulation can reduce adaptation^16^, but here we linked this effect directly with changes in excitatory-inhibitory balance. Dynamic frequency modulation did not reproduce these benefits. Even when lower-frequency hold periods improved offset encoding, network dynamics stayed close to fixed-frequency stimulation, with strong inhibitory recruitment and adaptation (Fig. 7G-J). This distinction suggests that reducing overall network drive through amplitude modulation differs from lowering stimulation frequency while maintaining continuous activation.

Closed-loop stimulation requires a measurable signal of the inhibitory network’s current state. The distinct dynamics of PV and SOM recruitment provide two potentially useful signals or mechanisms for this purpose. PV recruitment followed the onset transient and rose with amplitude and frequency (Fig. 2G), making it a readout of feedforward drive. SOM recruitment rose only once sustained drive passed 20 Hz, making it a readout of accumulating feedback inhibition. Limiting sustained amplitude while preserving transient amplitude reproduces the dynamic amplitude condition tested here, where excitatory and inhibitory populations did not differ detectably (Fig. 7B). Therapies that depend on inhibitory recruitment could select a pattern for the desired subpopulation. From our results, burst-type patterns (such as 10 Hz and 20 Hz Bursting) preferentially recruit PV interneurons, while TBS preferentially increases SOM activity over time.

The cortex is particularly sensitive to sustained stimulation intensity, which may determine inhibitory accumulation. PV neurons remained strongly responsive during onset and offset transients, but SOM neurons were not. Lower amplitudes of stimulation can reduce inhibitory activity and decrease adaptation, so that transient high amplitude bursts may be preferable for maintaining network excitatory-inhibitory balance or activating PV neurons transiently, while sustained stimulation results in strong inhibitory drive in the network. This work provides direct physiological insight into previous reports showing that dynamic or biomimetic amplitude modulation can improve percepts^50–52^.

### Distal facilitation is transsynaptic rather than direct

Facilitation of distal SOM neurons during 20-Hz stimulation and TBS is unlikely to arise from direct activation. Direct activation is confined within 40 to 60 µm of the electrode at comparable charge per phase^22^. Dense glutamate release stays within 20 µm of the site even at 15 and 20 µA^21^. Both bounds were measured under ketamine anesthesia, which broadens the spatial extent of cortical responses relative to wakefulness, so the direct-activation zone in awake recordings is unlikely to be larger than these estimates. Transsynaptic recruitment is therefore the most likely route to SOM neurons classified as increasing beyond 200 µm (Fig. 3D, Supp. Fig. 2C). Distance-binned intensity across the full pattern set supports the same reading, with every 20-µA pattern decreasing across distance except 20 Hz and TBS in SOM neurons, as well as 20 Hz and 10 Hz Burst in non-SOM neurons and 20 Hz and 200 Hz in PV neurons (Fig. 3E,G,H, Supp. Fig. 3H). Dynamic frequency patterns behaved like their fixed-frequency counterparts (Supp. Fig. 5). The temporal pattern of microstimulation can therefore produce more complex synaptic effects that depend on the specific stimulation pattern. A generic consequence of sustained stimulation would be expected to emerge across the fixed, burst, and biomimetic patterns as well, rather than being restricted to 20 Hz and TBS, which also has implications for the observed behavior of the VGAT-labeled population recorded previously^17^.

### Theta-burst stimulation can engage both inhibition and disinhibition

TBS facilitated and increased SOM activity within and across trials. Previous work has suggested that TBS can enhance inhibitory network activity through synaptic mechanisms^17,67,68^. However, recruiting more inhibitory interneurons does not necessarily reduce excitatory network output. SOM neurons can suppress PV interneurons, potentially producing disinhibition of excitatory neurons despite increased inhibitory interneuron activity overall^27,33,61^. Consistent with this possibility, SOM recruitment was stronger during TBS, and PV recruitment stronger during 20-Hz-Burst stimulation (Fig. 5C,D). Thus, the facilitating and depressing responses to TBS may arise from a complex balance between inhibition and disinhibition. Continuous high-frequency stimulation drives suppressive adaptation, while TBS may strengthen inhibitory recruitment gradually over time without immediate network collapse. Therapeutic benefits of TBS may therefore depend on SOM-mediated modulation of network plasticity as well as on inhibitory recruitment. Different microstimulation paradigms could selectively engage separate inhibitory circuit mechanisms according to the therapeutic or sensory goal.

### Depression during strong stimulation is a multi-cellular event rather than a metabolic limit

Rapid loss of activity during strong stimulation may be attributed to metabolic shortage, in which oxygen or glucose supply fails to meet demand. Several measurements instead point to activity-dependent membrane and synaptic mechanisms. Pericyte calcium falls during 40 and 100 Hz stimulation, and arteriole diameter near sphincter pericytes rises with frequency, so the cells that constrict vessels relax as frequency climbs^69^. Oxyhemoglobin also stays elevated through the depression that follows activation, while deoxyhemoglobin falls and the oxygen extraction fraction drops, so the tissue extracts a smaller fraction of the oxygen delivered^18^. Blood supply therefore rises across the same frequencies where network output falls (Fig. 2A,F). This rise indicates that the depression is not caused by a shortage of oxygen or glucose, since both increase at higher frequencies. A global shortage of delivered oxygen or glucose is therefore unlikely to drive depression, although these bulk measurements cannot rule out a local shortfall in oxygen delivery to individual neurons. Consistent with this observation, inhibitory neurons act on the suppressed population directly, with PV providing fast perisomatic inhibition and SOM accumulating over seconds within a pattern to suppress excitatory somatic activity once drive passes the threshold measured here.

Beyond the inhibitory circuit, several non-neuronal cell types converge on these same suppressed neurons. Pericytes and vascular smooth muscle relax as frequency climbs, uncoupling network output from blood supply^69^. Microglia act on a slower timescale, orienting their processes toward activated neurons during 10-Hz stimulation and preferentially contacting neurons whose activity has already depressed^23,70^. Microstimulation also drives activity in astrocytes primarily through neuronal signaling and suppressing or amplifying astrocyte calcium can increase or decrease stimulation-evoked neuronal activity, most strongly in the synaptically recruited neurons distal to the electrode^71,72^. Each cell type acts on the same suppressed neurons, layering onto the microcircuit near the electrode rather than reflecting a single metabolic limit. Calcium imaging alone cannot resolve whether astrocyte, microglial, and vascular activity deepen the depression or follow it.

### Limitations and future directions

Several limitations for this study should be considered. Differences in cortical state, area, and stimulation duration complicate comparisons with previous studies. Additionally, here we did not analyze sensory evoked activity but rather used a method to remove any large shifts in activity driven by window movement or sensory evoked-responses. Collecting multiple trials in multiple animals helps us account for variability in movement and sensory evoked activity. Future studies should evaluate how ongoing sensory experiences impact the responses to microstimulation. GCaMP7b lacks the temporal resolution to capture individual pulse responses^65,66^. Entrainment to individual pulses therefore cannot be measured here, and voltage indicators would resolve single-pulse dynamics at cellular resolution. New voltage indicators with increased spatial resolution can help understand the temporal dynamics more precisely^67^. Only somatic signals were analyzed, stimulation waveform asymmetry was not varied, and PV and SOM neurons were recorded in separate cohorts, limiting assessment of their interactions. The computational model provides a potential mechanistic framework but does not establish causality and cannot fully capture complex pattern-dependent or across-trial effects. Targeted manipulation of PV and SOM neurons is needed to validate their proposed roles. Future models should also incorporate other inhibitory populations, particularly vasoactive intestinal peptide (VIP) neurons^68^, which may contribute to disinhibition and across-trial changes in activity.

### Conclusion: temporal patterning shapes cortical recruitment and inhibitory stabilization

Collectively, these findings support a broader conceptual framework in which microstimulation-induced activity exists along a continuum from sparse, distributed recruitment to dense activation with increased inhibition depending on the stimulation intensity. Low-amplitude and transient stimulation patterns favor sparse recruitment, reduced inhibitory buildup, and sustained network responsiveness, potentially producing more naturalistic activity patterns. In contrast, strong continuous stimulation drives dense activation that progressively recruits inhibitory stabilization mechanisms, particularly SOM-mediated feedback inhibition, resulting in adaptation and suppression of cortical activity over time. The computational model reproduced the experimental results within this framework and explained the observed distance relationships, providing strong support for this understanding of circuit activation. This model demonstrated that inhibitory dynamics and their influence on network activation are likely tied to changes at the synapse, and direct stimulation bypasses dendritic integration of input which can result in unnatural continuous activation of recruited neurons.

These findings inform basic neuroscience, sensory neuroprosthetics, and therapeutic stimulation paradigms. From a basic neuroscience perspective, the present results provide further insight into excitatory-inhibitory circuit dynamics and support the idea that inhibitory neurons are preferentially recruited by strong cortical input. SOM neurons were weakly recruited at low amplitude and frequency, and their recruitment rose between above 40 Hz at 4 nC/phase. Time-to-peak separated the two populations beyond 175 µm from the electrode, placing PV recruitment in the feedforward path and SOM recruitment in the feedback path. In particular, the strong recruitment of SOM neurons during prolonged or intense stimulation suggests that these neurons may contribute substantially to activity-dependent suppression of cortical networks over time.

For sensory neuroprosthetic applications, stimulation strategies that minimize excessive recruitment of feedback inhibitory circuits may better preserve perceptual stability and reduce adaptation during prolonged microstimulation use. Dynamic amplitude modulation delivered brief high-amplitude transients on a low-amplitude hold, producing little adaptation within or across trials and reducing strong SOM activation. In therapeutic stimulation paradigms, these findings may also provide a mechanistic explanation for the reported advantages of theta-burst and other burst-based stimulation patterns over continuous fixed-frequency stimulation. The 20-Hz Burst pattern maximized PV recruitment while TBS raised SOM activity across trials, so a pattern can be chosen to maximize engagement of one inhibitory pattern rather than both. More broadly, understanding how distinct inhibitory subnetworks shape cortical responses to different temporal stimulation patterns may enable the development of stimulation paradigms that produce more effective and stable modulation of cortical activity while minimizing maladaptive network effects, such as excessive suppression.

## Methods

### Surgery, virus injection, and electrode implantation

Transgenic PV-Cre (n = 4) and SOM-Cre (n = 4) mice (Jackson Laboratories, Bar Harbor, ME) were used in these experiments. All procedures were performed on adult mice (> 4 weeks of age) housed in social groups under a 12 h light/dark cycle with ad libitum access to food and water.

Several weeks prior to experiments, mice received intracortical injections of AAV-Syn-GCaMP7 to enable calcium imaging of excitatory neurons and AAV-FLEX-tdTomato to label PV- or SOM-expressing interneurons, as previously described^17^. Injections were performed under aseptic conditions using a micromanipulator, glass pipette (5-10 µm tip diameter), and microinjector (Toohey Company, Fairfield, NJ, USA). Anesthesia was induced with ketamine (75 mg/kg) and xylazine (7 mg/kg) administered intraperitoneally (IP), with supplemental ketamine (40 mg/kg) provided as needed.

A rectangular craniotomy (∼4 mm per side) was made over the forelimb region of the primary somatosensory cortex (4 mm lateral, 1 mm anterior to bregma) using a dental drill. Viral injections were performed within this region using anatomical markers to inject around the forelimb somatosensory cortex, and the craniotomy was sealed with a glass cover slip and dental cement following the procedure. Mice were allowed to recover for at least two weeks before experiments. Prior to experiments, forelimb expression was assessed under a wide-field microscope by stimulating the forelimb with a cotton swab to confirm placement in forelimb cortex.

For the closed-window preparation, the cover glass was removed under isoflurane anesthesia, and a Michigan-style silicon microelectrode array (3 mm-long shanks, four 703 µm² sites per shank, 100 µm spacing; NeuroNexus Technologies, Ann Arbor, MI) was inserted into the cortex where there was clear viral expression. The array was secured with Kwik-Sil (World Precision Instruments) and resealed with a coverslip and dental cement. Following cessation of anesthesia, animals were monitored until awake, and recordings were initiated once normal behaviors (e.g., whisking, limb movement) were observed. In the open-window preparation, the coverslip was similarly removed and a single-shank Michigan-style array (3 mm shank, 16 sites, 703 µm² per site, 100 µm spacing) was inserted. When necessary, agarose was applied to stabilize the exposed brain and minimize motion artifacts.

In all cases, electrodes were implanted to a depth of 600 µm at a rate of 200 µm/s at a 30° angle using an oil-hydraulic microdrive (MO-81, Narishige, Japan). All animal care and procedures were performed with the approval of the University of Pittsburgh Institutional Animal Care and Use Committee and in accordance with regulations specified by the Division of Laboratory Animal Resources.

### Electrical stimulation

Electrical stimulation was delivered using a current-controlled TDT IZ2 stimulator controlled by an RZ5D processor (Tucker-Davis Technologies, Alachua, FL). Cathodic-leading, biphasic, asymmetric pulses (200 µs cathodic phase, 100 µs interphase interval, 400 µs anodic phase) were used, matching the waveform parameters from previous human studies^1,2,5,51^. This pulse shape was chosen because waveform asymmetry can influence neural recruitment^15,77–79^. Pulse amplitude controlled the charge per phase, while pulse frequency determined the number of pulses per second. Stimulation amplitudes were 10 and 20 µA (2 and 4 nC/phase) and frequencies ranged from 10-200 Hz. These parameters were selected to remain within established safety limits for both electrode integrity and tissue health^80–84^.

Stimulation was applied through a single electrode within the imaging plane at amplitudes of 10 µA and 20 µA (2 and 4 nC/phase). Each amplitude was tested with several temporal patterns, including 10-Hz, 20-Hz, 40-Hz, 100-Hz, and 200-Hz constant-frequency patterns; 10-Hz and 20-Hz Burst patterns; theta-burst stimulation (TBS); and biomimetic-amplitude or frequency-modulated waveforms (amplitude-modulated at 100 Hz or 200 Hz; frequency-modulated between 200-40-200 Hz or 100-20-100 Hz). In total, 11 stimulation patterns were presented at each of two amplitudes during every experimental session.

The order of stimulation patterns was pseudorandomized. Each pattern was delivered four times for 30 s, with 15-s inter-trial intervals to both increase sampling and assess the effects of repeated stimulation on neural activity. Generally, there was a ∼2-min break between each stimulus pattern delivery based on the time it took for the data to process.

### Two-photon imaging

A two-photon laser scanning microscope (Bruker, Madison, WI) and an OPO laser (Insight DS+; Spectra-Physics, Menlo Park, CA) tuned to a wavelength of 920 nm at 15 mW was used for the entirety of this study. The 16X 0.8 numerical aperture water immersion objective lens (Nikon Instruments, Melville, NY) with a 3-mm working distance. Imaging was in plane with the electrode site, 120-150 µm below the surface of the brain. To image neural dynamics during electrical stimulation, time series images were recorded at 30 Hz and 1.5× optical zoom. Time-series imaging was centered over the electrode contact so that the stimulating contact was visible during the imaging session, as described previously^16^.

### Image processing

All imaging data were preprocessed to minimize artifacts from motion and spontaneous sensory activity. Two main factors affected data quality: (1) brain movement relative to the imaging plane, particularly during open-window electrode insertions, and (2) neural responses to spontaneous or behaviorally driven sensory inputs (e.g., forelimb movement). Large movements or sensory evoked responses could raise the apparent intensity of certain patterns without reflecting real differences in stimulation-evoked activity, biasing the analysis. Therefore, we developed a preprocessing pipeline to account for these effects and isolate stimulation evoked-responses.

### Motion Correction and Frame Selection

Green (GCaMP7) and red (tdTomato) channels were loaded into MATLAB and converted from .*ome* to .*mat* format. Motion in the x-y plane was corrected using the NoRMCorre algorithm^85^, and corrected stacks were reconstructed as .*tif* files. The red channel, being calcium-insensitive, was used to identify residual z-motion.

Red-channel data were averaged across 20-frame bins (0.67 s per frame). Each binned frame was background-subtracted (Gaussian-filtered) and thresholded at three standard deviations above background to generate binary masks of neuronal structures. A stable “template” frame was selected, and all frames were compared to this template by pixelwise multiplication of binary masks to compute spatial overlap. Overlap values were normalized to the template’s self-correlation. Frames with normalized correlation < 0.5 were considered z-shifted and excluded from both channels. This threshold was selected based on visual validation that correlations above this threshold generally represented small shifts in the plane that still included the neurons of interest, while values below this threshold represented major shifts in the z-axis with different neurons in plane.

### ROI Extraction and Fluorescence Quantification

Motion-corrected, z-stable stacks were imported into ImageJ (NIH). Cells were manually outlined using a 2D projection of the pixelwise mean of fluorescence. Mean fluorescence over time was extracted for each ROI and analyzed in MATLAB (MathWorks, Natick, MA).

Fluorescence traces were temporally smoothed with a 15-point Gaussian filter and normalized to baseline to yield ΔF/F₀, with F₀ defined as the mean fluorescence during a 10 s pre-stimulation period.

### Separation of Stimulation-Evoked and Sensory-evoked or Spontaneous Activity

To isolate stimulation-locked responses from spontaneous or movement-related activity, fluorescence traces were decomposed using non-negative matrix factorization (NMF) implemented via the MATLAB *nnmf* function. NMF provides an additive, parts-based representation of neural population activity, constraining all components to be non-negative for biological interpretability. This method was selected over alternatives such as principal components analysis and independent components analysis, based on early pilot analysis in which it was found to better separate out stimulation induced responses from other non-stimulation related components.

We chose to use three components which were assumed to represent (1) stimulation-evoked responses, (2) sensory-driven activity, and (3) background or baseline fluctuations. This dimensionality was chosen based on pilot analyses showing that higher component numbers did not improve reconstruction accuracy or interpretability, while lower component numbers did not clearly separate noise from stimulation evoked-responses.

To improve interpretability and convergence, one temporal component was initialized with the known stimulus time course (we will call stimulation vector *S*). Specifically, a stimulus vector representing the timing and amplitude of the experimental stimulation was used to seed one row of the temporal component matrix (*H*). The corresponding spatial weight matrix (W) was initialized randomly. Thus, the decomposition took the form:

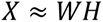

Where *X* represents the original response matrix (organized by neurons and timepoints), *W* represents neuron-specific weights for each component and *H* represents the temporal profiles of those components. NMF was solved using multiple replicates (10 iterations) to ensure stable solutions.

Temporal components were correlated with the stimulation vector *S* using Pearson’s r to identify those associated with the stimulation pattern. Non-stimulation components were excluded or subtracted, and the denoised activity was reconstructed as:

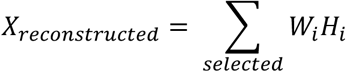

Slow baseline trends were fitted to non-stimulation periods and subtracted to correct for any gradual drift of the neural signal. Reconstructed traces were visually inspected to ensure retention of stimulation-evoked activity while minimizing spontaneous or sensory-evoked fluctuations.

### Quantification of Neural Activation

Cells were classified as active if their ΔF/F₀ during stimulation exceeded 3 standard deviations above baseline, as described previously^17^. For active cells, normalized fluorescence power was compared across stimulation parameters to identify preferential activation patterns.

### Separation of excitatory, inhibitory, and unclassified populations

The identification of tdTomato-expressing inhibitory neurons followed the approach described previously^17^. Briefly, neurons could not be reliably separated into tdTomato-positive and negative populations using a simple fluorescence intensity threshold, as red-channel intensity was distributed continuously rather than bimodally. Baseline fluorescence in the green (GCaMP) and red (tdTomato) channels exhibited a strong linear relationship, consistent with partial spectral overlap between the emission profiles of GCaMP and tdTomato. GCaMP fluorescence contributed to detectable signal in the red channel during stimulation, likely due to minor overlap in the emission spectra and the transmission ranges of the applied green and red filters.

To correct for this cross-channel contamination, we modeled the relationship between baseline green and red fluorescence using linear regression. Neurons with red intensity values greater than one root mean square error (RMSE) above the fitted line were classified as inhibitory, while neurons below the fit were classified as excitatory. Cells within one RMSE of the fit were considered unclassified, as their fluorescence could not be confidently attributed to either population. This regression-based classification method was previously validated and shown to perform comparably to more complex approaches^17^, providing reliable separation of inhibitory and excitatory populations while minimizing misclassification arising from spectral overlap.

All fluorescence data were baseline-normalized to account for any residual cross-channel influence, ensuring that the green-channel activity traces reflected true calcium-dependent dynamics.

### Data analysis and statistics

All data analysis and statistical analyses were performed in MATLAB (MathWorks, Natick, MA, USA) using custom-written scripts unless otherwise noted.

For temporal analyses, neuronal activity was quantified during predefined stimulation windows. “Onset” responses were calculated as the mean fluorescence during the first 3 s of stimulation, while “sustained” responses were calculated as the mean fluorescence during the final 10 s of the stimulation period unless otherwise stated. In analyses of dynamic amplitude modulation, onset and offset transient responses were additionally quantified within 1-s windows immediately following stimulus onset or offset transitions.

To quantify changes in activity over time during stimulation, a change index was calculated for each neuron as:

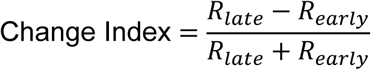

where *R_early_*was the mean activity during the first half of stimulation (0-15 s) and *R_late_*was the mean activity during the second half of stimulation (15-30 s). Positive values indicated facilitation or increasing activity over time, while negative values indicated adaptation or depression.

To characterize temporal response dynamics, neurons were classified into response-profile categories based on the magnitude and direction of activity changes throughout stimulation, similar to previous work^17^. Neurons showing strong rapid decreases in activity were classified as rapidly decreasing (RD), neurons with slower decreases were classified as slowly decreasing (SD), neurons with relatively stable activity were classified as non-decreasing/stable (ND), neurons with progressively increasing activity were classified as increasing (Inc), and neurons showing net suppression below baseline during stimulation were classified as depressed (Dep). Classification thresholds were determined from normalized temporal response profiles, comparing the first 1 s of activation to the later parts of the activation pattern, with criteria for different labels matching values reported previously for these classes^17^.

For frequency preference analyses, each neuron was assigned a preferred stimulation frequency corresponding to the stimulation condition that produced the largest mean response in the last 10 s. Frequency preference distributions were then compared across neuronal subtypes.

Spatial analyses were performed by calculating the Euclidean distance between each neuron centroid and the stimulating electrode location. For distance-based analyses, neurons were either analyzed continuously using regression-based approaches or grouped into spatial distance bins extending radially from the electrode tip. Mean activation intensity and temporal response classifications were compared across distance bins and neuronal subtypes.

Comparisons across stimulation frequencies, amplitudes, neuronal subtypes, temporal windows, stimulation patterns, trials, and spatial distance bins were performed using linear mixed-effects models (LME), with neuronal subtype, stimulation condition, or other experimental factors included as fixed effects and animal included as a random effect to account for non-independence of observations collected from the same animal. Model significance was assessed using analysis-of-variance tests of fixed effects. When multiple statistical comparisons were performed within a defined family of tests, p-values were adjusted using the Benjamini– Hochberg procedure to control the false discovery rate.

Regression-based analyses were used to examine trends across stimulation conditions. For analyses of responses across stimulation frequency, frequency was represented on a log_2_ scale. Linear mixed-effects models included neuronal subtype, log_2_ frequency, and their interaction, with animal included as a random effect. The interaction term was used to test if the relationship between neuronal subtypes and response changed across stimulation frequency.

For analyses of time-to-peak differences for the burst patterns, we analyzed the patterns individually and in combination. Cell-type differences were evaluated separately for each pattern using LMEs with neuronal subtype as a fixed effect and animal as a random effect. An additional LME was performed after pooling observations across all four patterns to test for an overall difference between neuronal subtypes.

Data are presented as mean ± SEM unless otherwise indicated. Statistical significance was defined as *p* < 0.05. Within figures, *= *p* < 0.05, ** = *p* < 0.01, and ***= *p* < 0.001.

### Computational model

A recurrent rate-based network model was developed to describe the dynamics of Exc, PV, and SOM neurons in response to microstimulation. Because we were trying to reproduce the population dynamics observed with calcium imaging, we chose to use a rate-based network model, where the activity of a population of neurons is treated as a continuous variable, rather than modeling individual action potentials (such as in a leaky integrate-and-fire model). The model incorporated recurrent excitation, reciprocal inhibitory interactions between Exc, PV, and SOM populations, and population-specific short-term synaptic dynamics. Specifically, PV neurons had strong initial input (the strength of which was determined by the stimulation parameters) while the SOM neurons had facilitating input (that increased as a function of excitatory neuron activity). Network activity was simulated using a forward Euler integration scheme with a time step of 1 ms.

Population activity was constrained to be non-negative using a rectified linear unit (ReLU) activation function. For each population, activity evolved according to first-order rate dynamics:

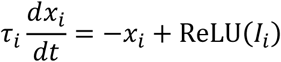

where *x_i_*represents the activity of population *i*, *I_i_*is its total synaptic and external input, and *τ_i_*is the corresponding population time constant. This form matches classic Wilson-Cowan formulation^86^.

The excitatory population received input according to:

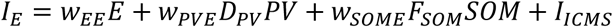

where *w_EE_*represents recurrent excitatory coupling, *w_PVE_*and *w_SOME_*represent PV- and SOM-mediated inhibition of E, respectively, and *I_ICMS_* represents the electrical stimulation input. *F_SOM_* and *D_PV_* were dynamic variables representing short-term facilitation or depression of the corresponding synaptic pathways.

PV activity was modeled as:

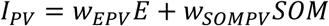

where *w_EPV_*represents excitatory recruitment of PV neurons by E, *w_SOMPV_*represents SOM-mediated inhibition of PV neurons, and *w_PVPV_* represents recurrent PV inhibition. SOM activity was modeled as:

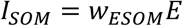

To capture activity-dependent changes in inhibitory and excitatory transmission, the model included dynamic facilitation of SOM-mediated inhibition and depression of PV-mediated inhibition. SOM facilitation was modeled as an activity-dependent process in which the target facilitation increased supralinearly with excitatory activity:

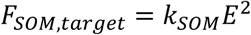

where *k_SOM_* controls the strength of the supralinear dependence. We made SOM facilitation supralinearly dependent on excitatory activity based on literature showing that Martinotti cells are supralinearly influenced by excitatory activity^43^. The facilitation variable evolved toward this target according to:

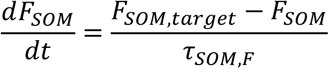

where *τ_SOM_*_,*F*_ determines the rate of facilitation. This form matches classic Tsodyks-Markram formulation^87^. A maximum value was imposed on *F_SOM_* to prevent unbounded growth.

PV-mediated synaptic efficacy was represented by a depression variable *D_PV_*, initialized at a baseline efficacy as a product of the stimulation parameters. During stimulation, the target value of *D_PV_* decreased as excitatory activity increased, such that stronger network activation produced greater depression of PV-mediated inhibition. The depression variable evolved according to:

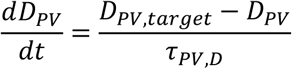

where *D_PV_*_,*target*_ was an activity-dependent function of E activity and *τ_PV_*_,*D*_ controlled the time course of depression. A lower bound was imposed on *D_PV_* to prevent complete elimination of PV synaptic efficacy. Thus, increasing excitatory drive resulted in progressively reduced PV-mediated inhibition, providing a dynamic negative-feedback mechanism analogous to short-term synaptic depression.

Microstimulation was represented as a pattern of instantaneous positive inputs delivered to the excitatory population. For conventional stimulation conditions, pulses were generated at the experimentally specified frequency and amplitude. Additional stimulation paradigms, including burst and biomimetic amplitude-modulated stimulation, were implemented by modifying the temporal structure and amplitude of the external input while maintaining the same underlying network dynamics. The model was simulated over an extended time window surrounding stimulation, with stimulation beginning 15 s after the start of the simulation. This provided a pre-stimulation baseline and post-stimulation period for comparison with experimental activity.

Because experimental measurements consisted of calcium fluorescence signals rather than instantaneous population activity, simulated Exc, PV, and SOM activity traces were transformed into calcium-like signals before comparison with experimental data based on previous reported latencies in calcium evoked activity^73,74^. Each simulated activity trace was convolved with a normalized temporal kernel consisting of a rapid rise component and slower exponential decay:

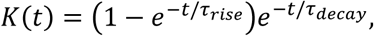

where *τ_rise_* = 30 ms and *τ_decay_* = 50 ms. The resulting convolved traces were used as the model-predicted calcium responses^88^.

Model parameters were semi-optimized using a random-search procedure. For each candidate parameter set, the model was simulated independently for each experimental stimulation condition. Each of the resulting Exc, PV, and SOM calcium-like traces were interpolated onto the temporal sampling grid of the experimental data. To preserve relative population amplitudes while eliminating differences in arbitrary model units, all simulated Exc, PV, and SOM traces across all stimulation conditions were jointly normalized using a single global maximum. Experimental traces were normalized using an analogous global normalization procedure. Performance was quantified using the mean squared error between simulated and experimental traces separately for Exc, PV, and SOM populations and across stimulation conditions. The total objective function was calculated as the sum of the mean losses for the three populations. Given we only ran a limited number of simulations (n<500) and the huge parameter space, the reported values may not be truly optimal, but reproduce the general shape of our experimental data and were adequate for this explanatory model. We stopped once we obtained parameters that satisfied the requirements visually.

### Spike deconvolution

Deconvolved spikes were not used in any statistical analyses and were provided solely for illustrative purposes. Calcium fluorescence traces were deconvolved using a custom greedy binary deconvolution algorithm implemented in MATLAB. Fluorescence was modeled as the convolution of a binary spike train with a single-exponential decay kernel. The imaging frequency was 30 Hz, the decay time constant (τ) was 0.2 s, and the kernel duration was set to 10τ. Spikes were evaluated sequentially at each time point and retained if their inclusion reduced the sum of squared errors between the observed fluorescence trace and the reconstructed signal. Thus, at each time point, a spike was retained when its inclusion resulted in a closer match between the modeled and observed fluorescence signals.

## Acknowledgments

This work was supported by NIH NINDS R01NS105691, NIH NINDS R01NS115707, NIH NINDS R01NS129632, NSF CAREER CBET 1943906, the University of Pittsburgh Senior Vice Chancellor for Research’s Pitt Momentum Scaling Fund, and The Pittsburgh Integrated Translational Neuroscience Institute Funds. C.H. was supported by the NINDS T32 NS08674932 training award for the study of the neurobiology of neurological disease and by the NIH BRAIN Initiative F32MH130022.

## Author contributions

C.H.: conceptualization, methodology, software, formal analysis, investigation, data curation, visualization, funding acquisition, writing of the original draft. T.K.: conceptualization, methodology, resources, supervision, project administration, funding acquisition, review and editing.

## Competing interests

C.H. is employed by Blackrock Neurotech, a company developing technology related to the work reported here. Data were collected at the University of Pittsburgh before that employment began, and the employment did not influence the content or conclusions of this study. Blackrock Neurotech had no role in study design, data collection, analysis, or the decision to publish. T.K. declares no competing interests.

## Data and code availability

Data and code will be made available upon full publication.

## Ethics approval

All animal care and procedures were performed with the approval of the University of Pittsburgh Institutional Animal Care and Use Committee and in accordance with regulations specified by the Division of Laboratory Animal Resources.

---

**Supplementary Figure 1:**
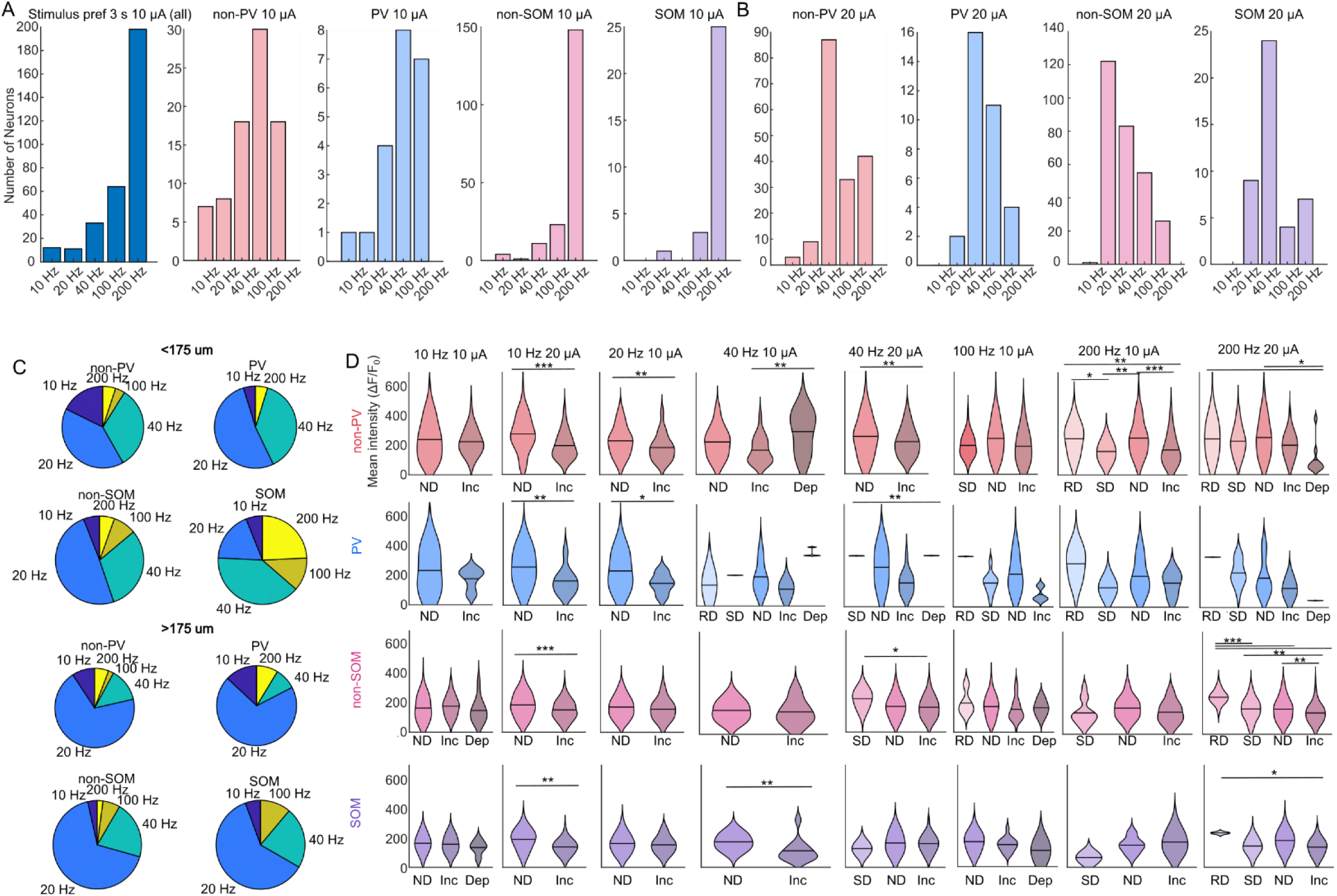
Amplitude reverses the preferred stimulation frequency, and distance shifts the SOM preference toward higher frequencies and neurons generally towards depression. A) Stimulus frequency preferences in the first 3 s across each cell type and combined across all cell types for 10-µA patterns. At low amplitudes, higher frequencies tend to drive consistently stronger responses so that most neurons have a preference for 100 or 200 Hz. B) Stimulus frequency preferences in the first 3 s across cell type for the 20-µA pattern. At higher amplitudes, the depression caused even within the first few seconds results in weaker activation by the higher frequencies, so that most neurons had the strongest responses for the 20 or 40 Hz patterns. C) Frequency preferences in the last 10 s of microstimulation for neurons less than and greater than 175 µm from the electrode. While most neurons have the largest preference for 20 Hz in the first trial, SOM neurons that are within 175 µm of the electrode are much more likely to have a preference for the higher frequency regimes (40, 100, and 200 Hz). D) Violin plots of the average distance of neurons based on the classification of their temporal response pattern. Consistently, “Increasing” neurons are closer to the electrode than “Non-decreasing” neurons, and the one exception is 20 Hz at 20 µA in SOM neurons (Fig. 3D). Neurons that have decreases during stimulation (RD or SD) primarily occur at the higher frequency regimes, which drive fast decreases in activation shortly after stimulation onset.

**Supplementary Figure 2:**
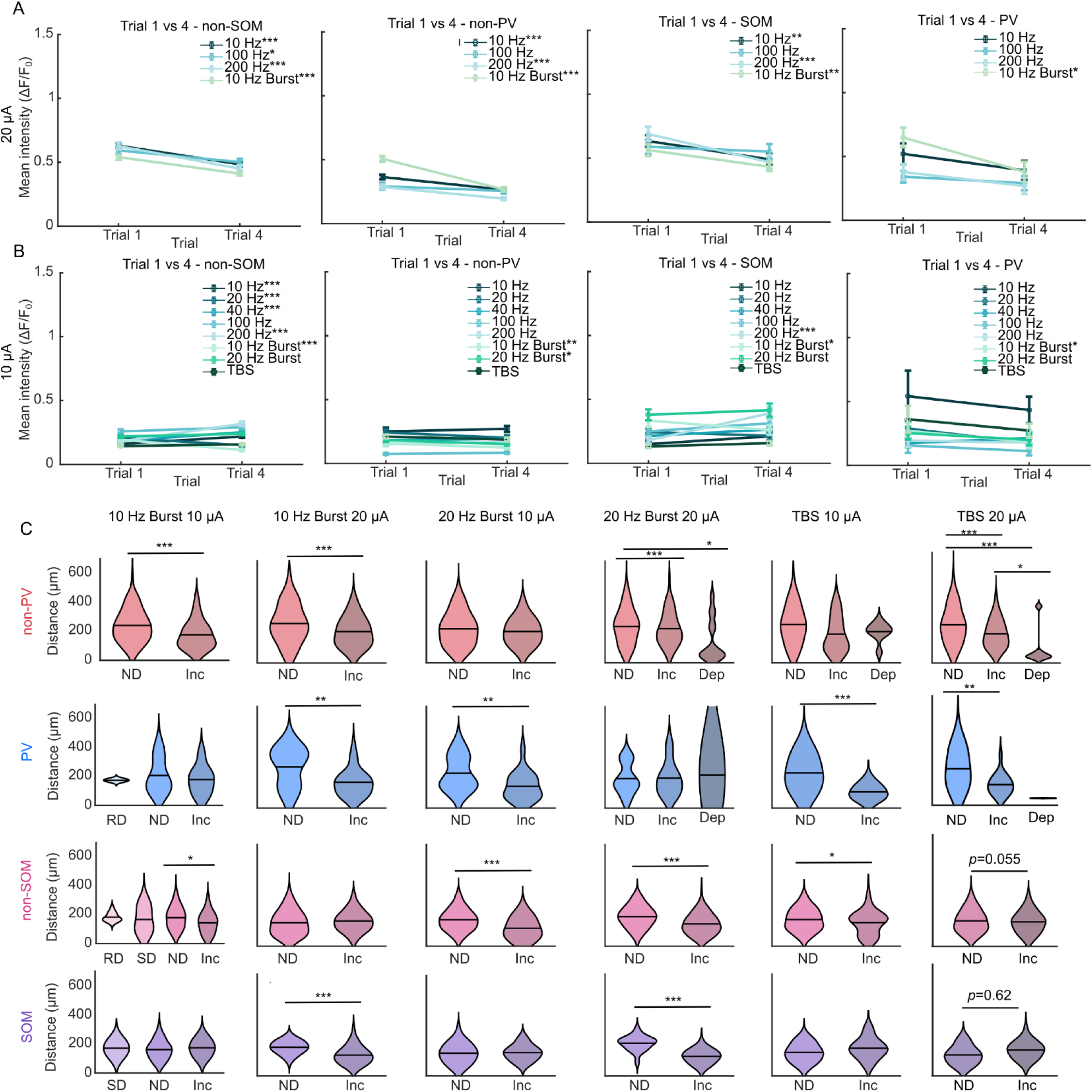
Lower amplitudes produce less adaptation across trials and burst patterns, while all amplitudes produce increasing responses closer to the electrode except for TBS in SOM neurons. A-B) Adaptation of evoked intensity from trial 1 to 4 for 20 µA (A) and 10 µA (B) patterns. The higher amplitude results in decreases in adaptation across trials, while the low amplitude does not result in as much decrease, and at times results in facilitation across trials, supporting that low amplitudes may be better for maintaining or increasing activity over time. C) Violin plots of the average distance for different classes of neurons. Similar to the fixed frequency patterns, ND neurons tended to be farther from the electrode than Inc neurons, except in SOM neurons when stimulated with TBS. SOM neurons had “increasing” neurons farther from the electrode relative to non-adapting neurons for both TBS at 10 µA and 20 µA, which is the opposite of what is observed for the other cell types, although these effects did not reach significance (*p* > 0.05).

**Supplementary Figure 3:**
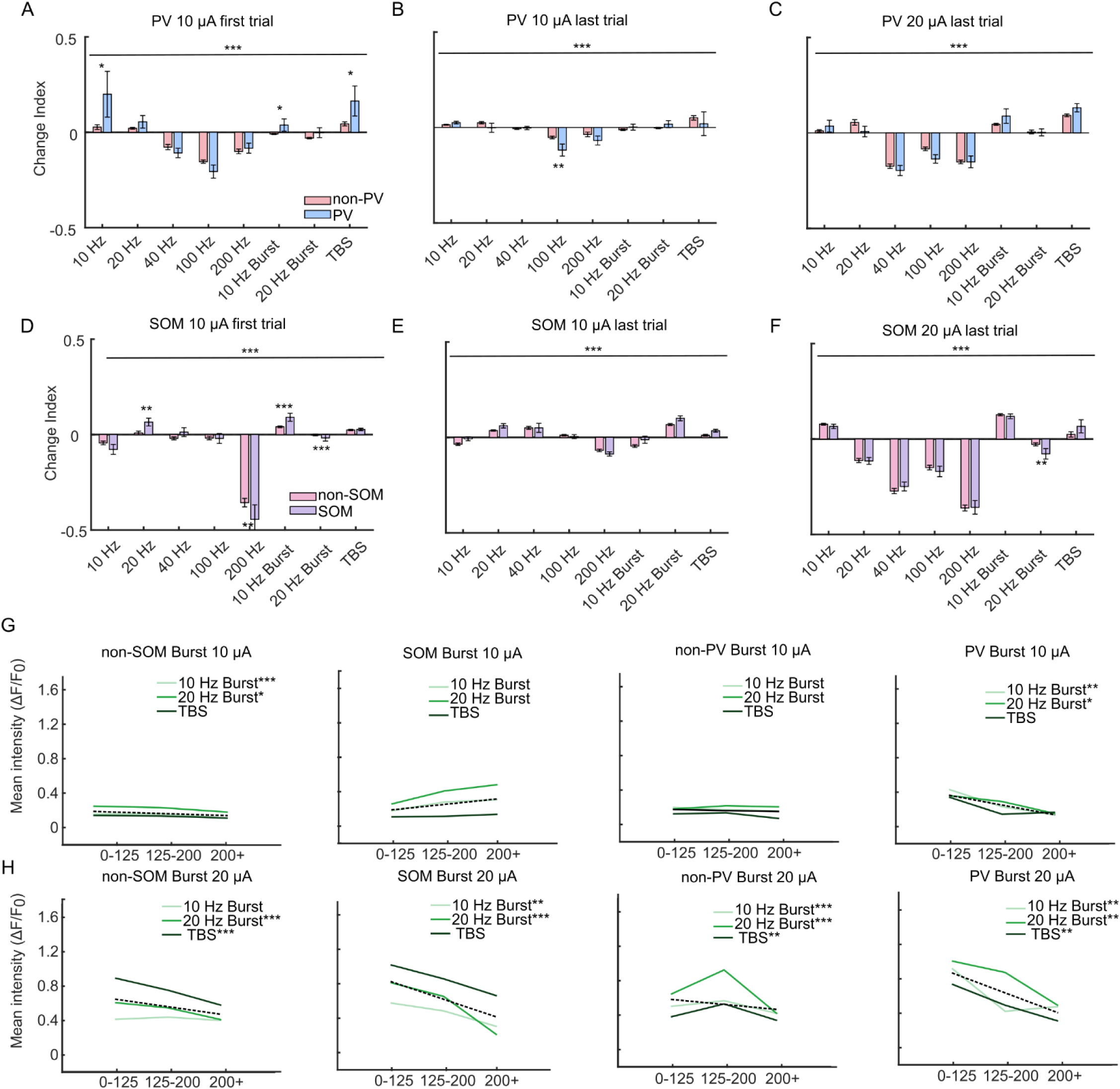
Burst patterns produce more facilitating responses within trial, especially in the last trial. Higher amplitude burst patterns consistently drive stronger responses near the electrode. A-F) Bar plots showing the change indices for different stimulation patterns for PV (A-C) and SOM (D-F) animals. Generally, the change indices were smaller (in both directions) for lower amplitudes and later trials, likely because the network has already showed some long-term adaptation preventing achievement of higher intensities and concomitant adaptation. (E-F). Burst patterns were generally better at driving facilitation, especially in the last trial except for SOM with the 20-Hz-Burst pattern which drove adaptation. G-H) Line plots showing the change in intensity across bins for 10- µA patterns (G) and 20-µA patterns (H). Similar to the fixed frequency patterns, 10-µA patterns tended to produce much less differences across bins due to the lower overall intensity of activation, although there were still some significant decreases across distance for 10 and 20 Hz Burst (*p <* 0.05), but not for the TBS pattern. At 20 µA, all patterns had significant changes across distance, except for TBS with SOM neurons and 10 Hz Burst 20 µA with non-SOM neurons.

**Supplementary Figure 4:**
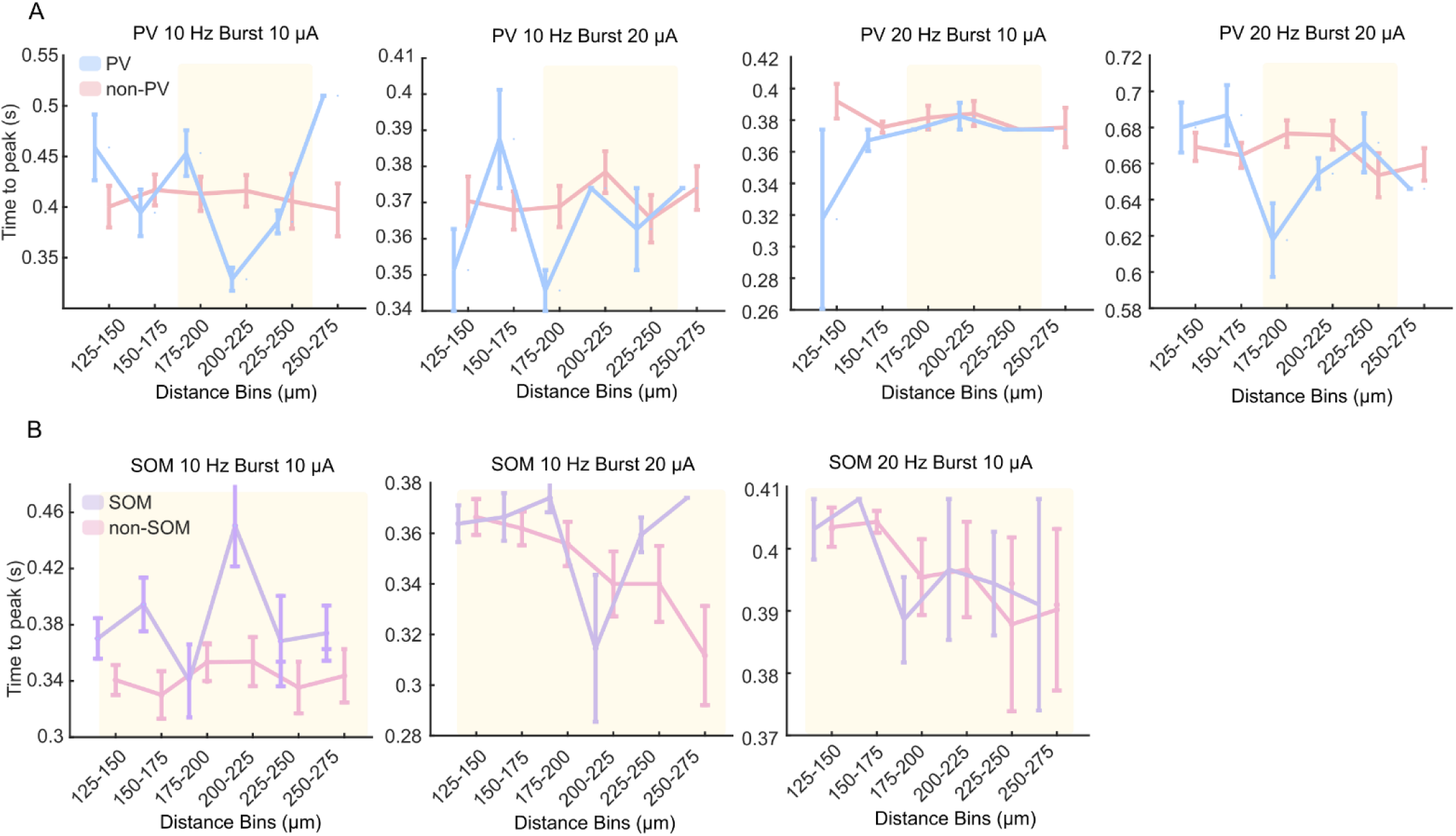
PV neurons peak at or before non-PV neurons and SOM neurons peak later than non-SOM neurons across distance. A-B) Line plots showing the average time to peak in different distance bins for PV (A) and SOM (B) animals. Error bars show the standard error across neurons. The highlighted regions show the distance bins with the maximal differences and were used for evaluation of group differences (175-250 µm for PV neurons and 125-275 µm for SOM neurons). Generally, PV neurons had faster or similar times to peak compared to non-PV neurons, while SOM neurons tended to have slower times to peak compared to non-SOM neurons. Note that SOM 20 Hz Burst 20 µA was not included here because it is included in the main figure 6A.

**Supplementary Figure 5:**
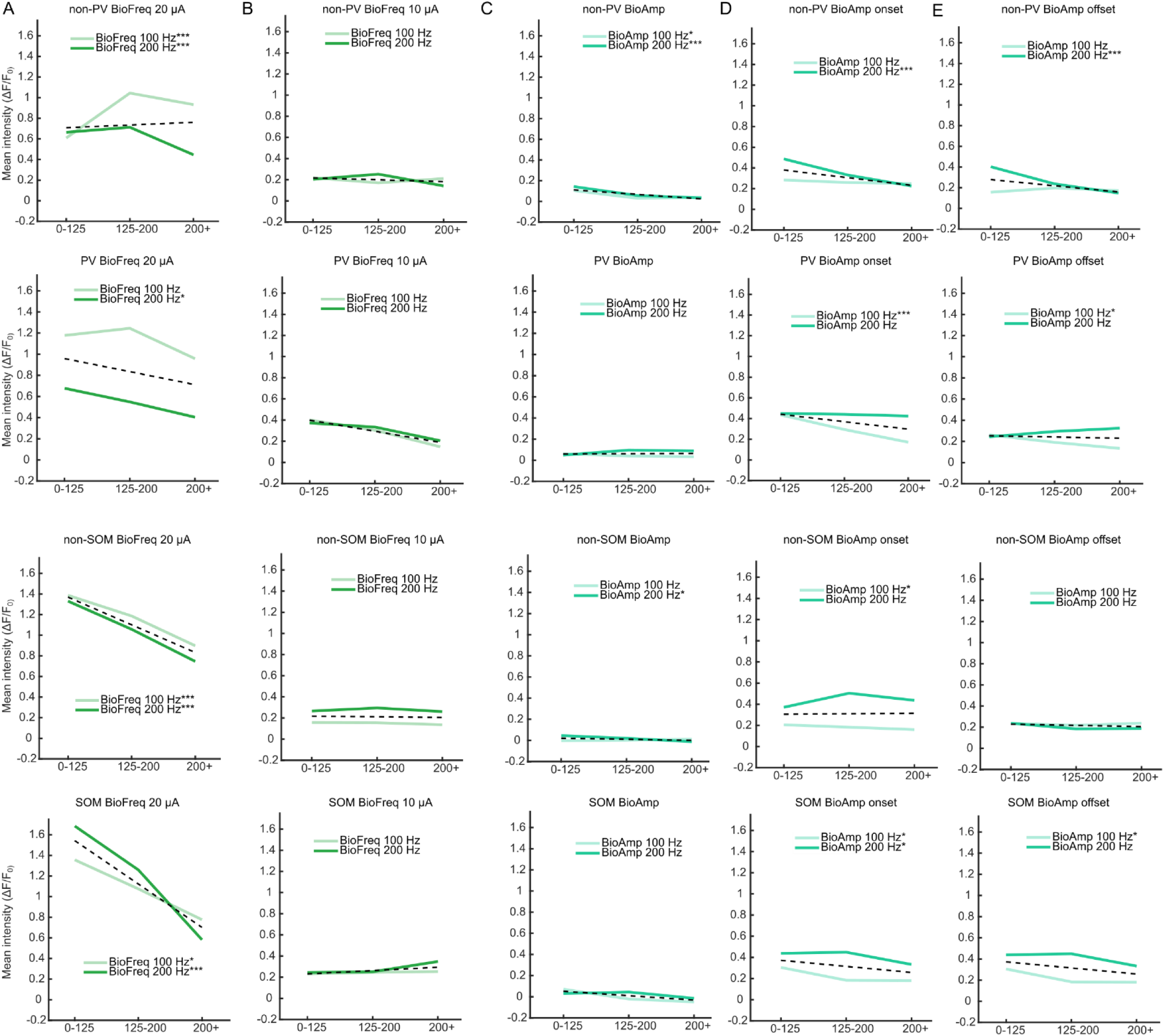
Dynamic frequency patterns lose intensity across distance like fixed-frequency patterns, while dynamic amplitude patterns change much less. A-E) Line plots showing the change in intensity across bins for the different dynamic patterns (A-C) as well as highlighting the onset and offset periods for the dynamic amplitude modulated pattern (D-E). Similar to the fixed frequency and burst patterns, the dynamic frequency patterns generally resulted in decreases in intensity across distance bins except for non-PV neurons with the BioFreq-100 Hz, 20-µA pattern, which showed some increase from the 0-125 bin to the 125-200 bin. The 10-µA patterns again did not drive much change across distance, with none of the patterns reaching significance. Across the whole pattern the dynamic amplitude patterns changed little with distance except in non-PV neurons, where both patterns decreased significantly. When focusing just on the high amplitude transients in which the strongest activity is driven, we see that there are some decreases across distance, but these trends are much weaker than the other patterns and most relationships are not significant.

